# CDR3 binding chemistry controls TCR V-domain rotational probability and germline CDR2 scanning of polymorphic MHC

**DOI:** 10.1101/796052

**Authors:** Joseph S. Murray

## Abstract

The mechanism which adapts the T-cell antigen receptor (TCR) within a given major histocompatibility complex (*MHC/HLA*) genotype is essential for protection against pathogens. Historically attributed to relative affinity, genetically vast TCRs are surprisingly focused towards a micromolar affinity for their respective peptide (p) plus MHC (pMHC) ligands. Thus, the somatic diversity of the TCR with respect to MHC-restriction, and (ultimately) to pathogens, remains enigmatic. Here, we derive a triple integral solution (from fixed geometry) for any given V domain in TCR bound to pMHC. Solved complexes involving HLA-DR and HLA-DQ, where genetic linkage to the TCR is most profound, were examined in detail. Certain V domains displayed rare geometry within this panel—specifying a restricted rotational probability/volumetric density (*dV*). Remarkably, hydrogen (H) bond charge-relays distinguished these structures from the others; suggesting that CDR3 binding chemistry dictates CDR2 contacts on the opposite MHC-II alpha helix. Together, these data suggest that TCR recapitulate *dV* and specialise target pMHC recognition.

## Introduction

**T**-cell antigen receptors (TCR) and antibodies (sIg) are individualized within each precursor of a given T-cell or B-cell clone. TCR (α/β type) have genetically variable (V) domains, wherein *complementary determining regions* (CDR1⍰CDR3) contain closest amino acid (a.a.) contacts with the peptide (p) plus *major histocompatibility complex* (MHC) protein (together abbrev., pMHC) composite ligand. CDR1 and CDR2 are encoded in the germline, via the particular V-region DNA segment involved in the *RAG1*/*RAG2* recombination mechanism responsible for somatic construction, together with the *D*- and/or *J*-segment(s), of the third, CDR3 [1⍰5]. This paper seeks to understand two puzzles of the TCR-pMHC interaction, wherein a novel examination of the first can be used to re-examine the second in the context of existing evidence. Firstly, compared to antibodies and indeed other protein:protein binding reactions, TCRs display quite low (μM) binding affinities for pMHC ligands [6⍰8]. Nevertheless, like antibodies, TCRs have exquisite specificity, where single peptide or MHC a.a. changes can dictate T-cell reactivity or non-reactivity [9⍰17]. Second, the nature of T-cell selection in the thymus prior to seeding secondary lymphoid tissues is thought to operate on the basis of this same relative TCR:pMHC affinity, where overt ‘self’ peptide plus self-MHC recognition (*negative selection*) leads to deletion from the repertoire; as does a complete lack of recognition (*positive selection*) [4]. This intermediate TCR:pMHC binding in the fetal thymus forms the basis of ‘simultaneous’ self plus pathogen recognition in adulthood, *i*.*e*., the *MHC-restricted* adaptive immune response [5, 18]. Somatic CDR3 loops are roughly positioned atop the peptide component in solved TCR:pMHC complexes [9⍰18]; although more recent investigations have challenged the notion that the most diverse component of the ligand (the peptide) is particularly interfaced with the most diverse regions of the receptor (CDR3α/β) [12, 19].

For example, we previously reported that one of the CDR3 loops (CDR3α) also makes consistently close contacts with a conserved HLA-A component, *i*.*e*., the *R65-motif* (a.a., R65-X-X-K68-A69-X-S71-Q72) [20]. *Burrows et al*., also found conserved CDR3 contacts directly with MHC-I [21]. More recently, *Sharon et al*., formally linked germline TCR usage to HLA, particularly via the class-II alleles [18]. Here, the *STCR-Dab* (www.opig.stats.ox.ac.uk/webapps/stcrdab/) was used to identify all current TCR complexes involving HLA-DR and HLA-DQ (3.0 Å resolution cut-off). Note, also included are *PDB* 4H1L, at 3.3 Å, and 3T0E, at 4.0 Å (see below). A simple relationship between the observed (*i*.*e*., measured) CDR3⍰CDR2 “pitch” angle for a given V domain and its predicted (*i*.*e*., calculated) CDR3⍰CDR2 pitch angle was observed [20]. Here, a linear relationship between calculated pitch and a new measure, “*dV*”—by multivariable calculus, was found. As is discussed, the simple volume element, *dV*, interprets V-domain orientation into a rotational probability involving an apparent CDR211MHC α-helix scanning function (*d*θ). While the *dV* was unique for each TCR on each pHLA-DR, one TCR displayed a dramatic restriction in *d*V for Vβ. This was isolated to a charge-relay H-bonding mechanism for CDR3β; hence, the chemistry of “somatic-TCR” dictates positioning of “germline-TCR” (see **Discussion**). Within the seven pHLA-DQ structures, the two *highly-restricted dV* TCR (*PDB* 4OZG & 4OZH) displayed distinct, yet functionally similar mechanisms that shifted the same TCR:MHC H-bond by one MHC a.a. position (relative to the nominal *dV* structure, 4OZF). This involves an additional H-bond (MHC:MHC) also found in suitable *charge-relay* mechanisms of 4OZG and 4OZH.

## Results

*PDB* files of solved TCR:pHLA-DR and TCR:pHLA-DQ structures were used to investigate V-domain geometry amongst the available complexes, *i*.*e*., involving similar (but different) TCR, and/or similar (but different) pMHC. All of these structures share the canonical (diagonal) orientation of the TCR over pMHC, which was one of the earliest observed similarities between different complexes [13]. As indicated in the summary tables (Tables 1A⍰1D; separation into different tables is simply for clarity) all sequences for eaf helix, then the angle at D66 to the ch component of all of these structures are available under the appropriate *PDB* file name at the *NCBI* (www.ncbi.nlm.nih.gov).

**Table 1A.**
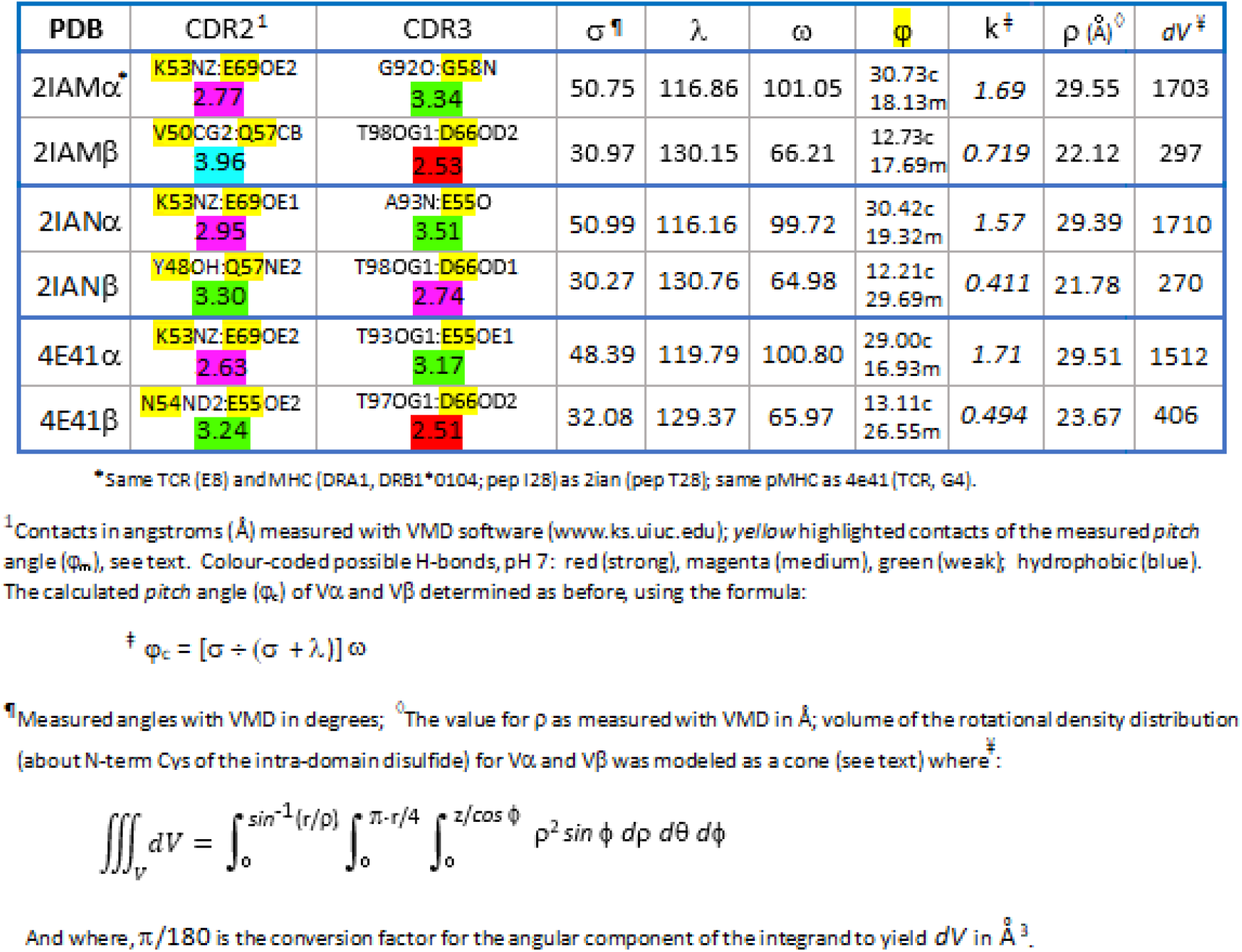
Summary Geometry TCR:pHLA-DR

### TCR-pMHC Geometry

Shown in Figures 1A and 1B is an example of the geometry analysis based on fixed a.a. positions in the HLA-DR/HLA-DQ grooves (*e*.*g*., *PDB* 1J8H). We originally used the concept of *Euler* angles (www.mathword.wolfram.com/EulerAngles.html) to establish the basic method (for a TCR-Vα cohort in pHLA-A2 complexes) [20]; here, the method was modified for the available solved TCR-pMHC-II structures (summarized in Tables 1A⍰1D). In brief, the analysis is based on measuring three angles corresponding to the *twist* (ω), *tilt* (λ), and *sway* (σ) of each V domain over the pMHC.

**Figure 1.**
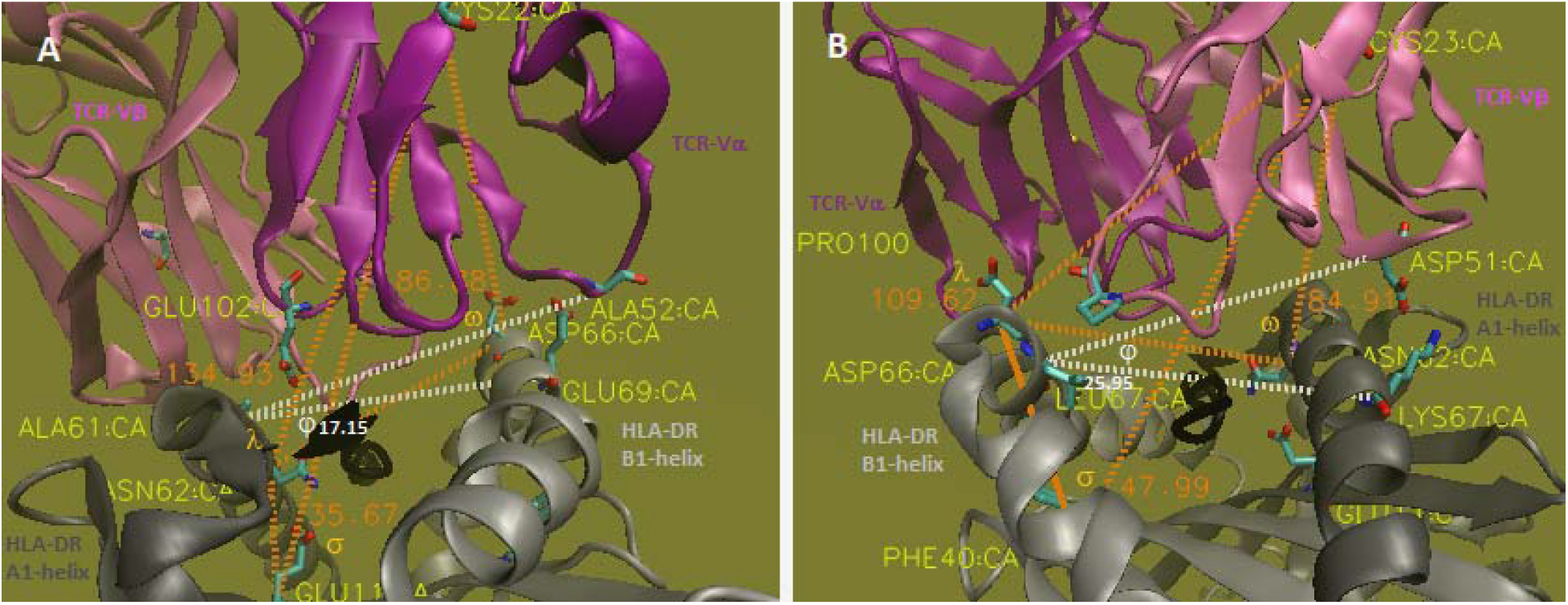
The twist/tilt/sway of TCR-Vα and -Vβ relative to pMHC-II (A & B, respectively). Illustrated by the example of 1J8H. This same analysis was performed on all 19 structures (Tables 1A ⍰1D). All angle measurements were from C*α* in *VMD-1*.*9*.*1* (www.ks.uiuc.edu) used to examine the diversity through the 38 V-domains: (i) in-plane to the MHC-groove {*twist* = *ω*}, with (ii) displacements perpendicular to the groove {*tilt* = *λ*}, including (iii) side-to-side variation {*sway* = σ}. The a.a. positions used as coordinates for angular measures (*dotted orange* vectors) across structures were fixed and are labelled (see text). The measured incline of a V domain, or “pitch” {*pitch* = *φ*_m_}, is shown by *white dotted* vectors and could be approximated by the equation: *φ*_c_ = [σ ÷ (*λ* + σ)]*ω* (see text, Tables 1A⍰1D). Side chains of a.a. in measurements are shown by *CPK licorice*; C*α*-backbones are in *new cartoon* and labelled; peptides are in *black*. Note from A to B the view of the structure rotates 180°. Figures are the original output of the *PDB* file as analysed in *VMD-1*.*9*.*1*. Best viewed at 100%.

For HLA-DR a vector from the DRA a.a. N62 alpha-carbon (N62:Cα) to DRB1/3/5 a.a. D66 (D66:Cα; Cα used for measurements unless otherwise noted) bisects the MHC-groove from the DRA α-helix to the DRB1/3/5 α-helix, then the angle at D66 to the Vα central cysteine (C22) is computed with the *VMD angle-label* tool. This ω-angle can be seen in Figure 1A (Vα) and 1B (Vβ; where the vectors run D66:N62:C23) as dotted *orange* lines near the *melon*-coloured “ω” symbol, 86.88° (Vα), 84.91° (Vβ). The E11 (beta-sheet floor) to N62 to C22 (*tilt*-angle) is similarly shown (1A) near the “λ” symbol, 134.93°; for Vβ, D66 is used (1B), F40:D66:C23, at 109.62°. Finally the σ-angle is measured from N62:E11:C22 (Vα) or D66:F40:C23 (Vβ), here at 35.67° and 47.99°, respectively. For HLA-DQ TCR:pMHC structures, Vα *twist* is based on a DQA a.a. N62:Cα to DQB D66:Cα vector bisecting the groove, where the angle vertex is at D66 to Vα C23:Cα. Here, N11:Cα (not E11 found in DR) to the N62 vertex to C23 defines *tilt*, and the reverse N62:N11:C23 is *sway*. For the DQ Vβ’s, angle a.a. are the same as in DR structures. Specifically, *twist* is defined by D66:N62:C23, *tilt* by F40:D66:C23, and *sway* as D66:F40:C23 (Table 1D). We formulated the equation:

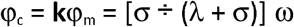

that predicts the pitch of a given V domain (*pitch* = angle φ) from the ω, λ, and σ (orientation). By contrast, measuring the φ-angle is based on finding the closest contact between the domain CDR2 and the α-helix that is opposite CDR3 across-the-groove, N62 (for Vα), or D66 (for Vβ) (*yellow* highlighted in Tables 1A⍰1D). The vectors are directed from that CDR2 a.a. to the across-the-groove α-helix a.a. in closest contact with CDR3, then back to the α-helix a.a. that is in contact with said CDR2 a.a. (*white* dotted lines, Figs. 1A & 1B). In Tables 1A⍰1D the calculated (φ_c_) and measured (φ_m_) pitch angles are shown along with the **k** values. Thus, to the extent k approaches 1.00 indicates correlation between the measured pitch and calculated pitch values. ***Statistics***: HLA-DR group: *mean k* = 0.99 ± 0.36 (*s*), *n* = 24, *t* = 0.915, *μ*_0_ = 1.00, *p* = 0.376, 95% CI, 0.880⍰1.10; (paired *Student*’s *t*-test; www.graphpad.com/quickcalcs; www.select-statistics.co.uk). HLA-DQ group: *mean k* = 0.88 ± 0.35 (s), *n* = 14, *t* = 1.24, *μ*_0_ = 1.00, *p* = 0.234, 95% CI, 0.750⍰1.01. Overall (DR & DQ): *mean k* = 0.95 ± 0.35 (s), *n* = 38, *t* = 0.85, *μ*_0_ = 1.00, *p* = 0.402, 95% CI, 0.870⍰1.03.

### Triple Integrals

We observed that the closest CDR2α contacts with the α2-helix ‘across-the-groove’ from the conserved R65 contact with CDR3α in HLA-A2 structures specified a polymorphic region from a.a. H/R151 to a.a. A158 [20]. Similarly, the CDR2α closest contacts range for the DR structures here (Tables 1A⍰1C) implicate a polymorphic DRB α-helix range from a.a. E69 to a.a. T77; for CDR2β, a DRA α-helix contacts range: a.a. Q57 to a.a. K67. For the DQ complexes: CDR2α contacts range is the polymorphic DQB α-helix a.a. E69 to a.a. D76; and for CDR2β, DQA: Q57 to H68, is also polymorphic (Table 1D). Thus, the hypothesis that if these α-helical a.a. are indeed swept/scanned by CDR2 of the V-domain, that one should be able to model such with integration from spherical coordinates in each structure [22, 23]. To normalize the equation across structures and use defined geometry, the central cysteine (C22/23) was chosen as a point rotating from the fixed a.a. position that defines *twist, tilt*, and *sway* (Figs. 1 & 2). As shown in Figure 2A & 2B, the coordinates model a ‘slice’ of a cone, where for each of the 38 V-domains a measured distance was taken between the C22/23 and said α-helix a.a. (*e*.*g*., 2IAM Vβ measures 22.12 Å)—this is the *rho* (ρ) distance in the equation. The other measure was the angle ϕ, which is simply the difference from 180° of the previously determined *tilt* angle (λ). The other variables were derived by trigonometry (see Fig. 2B).

**Figure 2.**
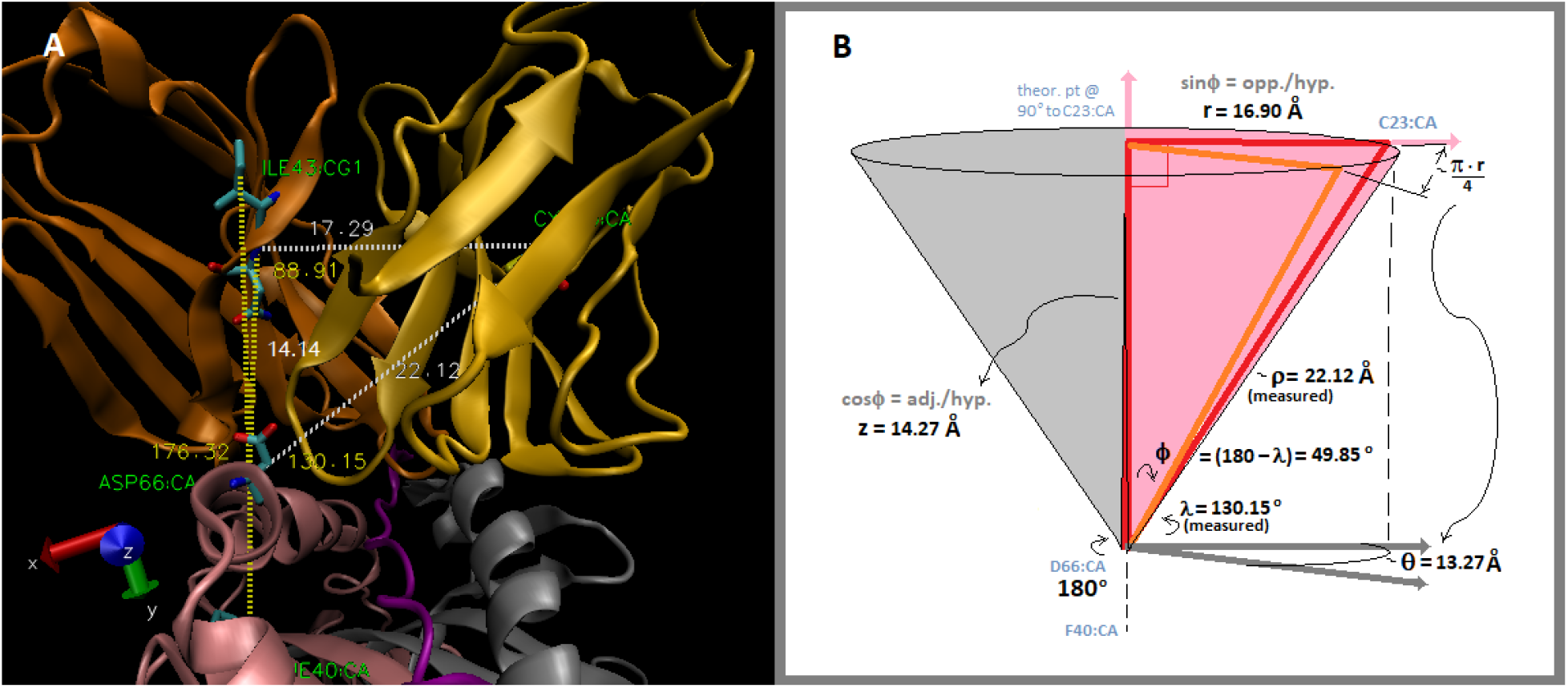
Derivation of the V-domain rotational volumetric-density equation. The same a.a. positions used for the tilt (*λ*) angles were used to define the phi (*ϕ*) angles (*e*.*g*., 49.85° for this structure, 2IAM V*β, gold*) by subtraction of *λ* from a theoretical 180° vector. This is illustrated with actual positions in 2IAM (A), showing the 180° approximating, 176.32° vector; the ∼ 90°, 88.91° angle; the ∼ 14.27 Å, 14.14 Å line segment; and the ∼ 16.90 Å, 17.29 Å line segment, *i*.*e*., to show the essential geometry within a structure; these are more accurately calculated by trigonometry in the derivation (B). Additionally, the rho (*ρ*) segment from the appropriate *α*-helix position (see *λ* definitions) to a given V-domain’s C22/23 was measured (here, 22.12 Å). The cone (full rotation) includes a probability (slice volume) through a *scanning path* (*dθ*) for CDR2. Figure (A) is the original output of the *PDB* file as analysed in *VMD-1*.*9*.*1*. Figure (B) is a diagram constructed in *MS-3D-paint*. Best viewed at 100%.

This is integration of a volume element in spherical coordinates (all *dV* values in Tables 1A⍰1D). As discussed above it is not surprising that the (π · r / 4) values were close to the actual distances between CDR2 contacts in the aforementioned MHC α-helix ranges using this approach, *e*.*g*., 13.27 Å, Figure 2B. Thus, the volume of the cone slice for each V-domain was determined upon integration for the three spherical coordinates, rho (ρ), theta (θ) and phi (ϕ), where the upper limit of each integral is derived from the measured ρ and ϕ values of each V domain as shown:

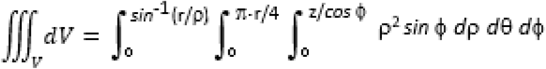

Since ρ and ϕ are measured for each V domain, derivations of upper limits for the first and third integrals simply provide trigonometric relationships.

Two example solutions are given below (parameters as defined, Figs. 1 & 2; text)—*for all 38 triple integral solutions*, see **Suppl.1.I**\*. The (π · r / 4) circumference segment is used as the upper limit of the *d*θ integral because it is accurate to a *path, i*.*e*., a distance; formally (for θ-angle in degrees): π · r (θ) / 180 = *arc length* in Å [22]. Hence, *d*ϕ is the only integrand for conversion (multiplying by π ÷ 180) to yield the *cubic angstrom* (Å^3^) unit of volume. The C22/23 cysteine is historically used as the center of any given V domain [10] and each V-domain cone-slice is thus the volumetric-density through its CDR2 scanning path (*e*.*g*., Fig. 2B). The calculus interprets a comprehensive geometry of the V domain into a probability of scanning using only the ground-state structure, *i*.*e*., without crystallographic data on theoretical conformers [24].

**Table 1B.** Summary Geometry TCR:pHLA-DR

| PDB | CDR2<sup>1</sup> | CDR3 | $\sigma^{\dagger}$ | $\lambda$ | $\omega$ | $\phi$ | $k^{\ddagger}$ | $\rho(\text{\AA})^{\diamond}$ | $dV^{\#}$ |
| --- | --- | --- | --- | --- | --- | --- | --- | --- | --- |
| 1J8H$\alpha^{\dagger}$ | A52CB:E69CD<br>3.54 | E102CD:A61CB<br>4.07 | 35.67 | 134.93 | 86.88 | 18.17c<br>17.15m | 1.06 | 29.77 | 740 |
| 1J8Hβ | D51OD1:K67NZ<br>2.54 | P100CG:L67CD1<br>3.43 | 47.99 | 109.62 | 84.91 | 25.85c<br>25.95m | 0.996 | 27.91 | 1734 |
| 1FYTα | A52CB:E69CD<br>4.42 | K103NZ:Q57OE1<br>4.36 | 37.11 | 133.00 | 86.29 | 18.81c<br>16.90m | 1.11 | 29.70 | 826 |
| 1FYTβ | D51OD1:K67NZ<br>2.67 | P100CB:D66CB<br>4.96 | 47.87 | 109.97 | 85.82 | 26.03c<br>23.44m | 1.11 | 28.04 | 1747 |
| 3TOEα | L50CD1:T106CG2<br>3.65 | K96NZ:Q57OE1<br>4.06 | 38.06 | 132.31 | 88.95 | 19.87c<br>21.36m | 0.930 | 29.99 | 855 |
| 3TOE$\beta^{\ddagger}$ | E50OE1:K67NZ<br>3.77 | N100CB:Q99CD<br>4.08 | 44.41 | 110.49 | 76.75 | 22.00c<br>20.98m | 1.05 | 24.63 | 1024 |
| 4H1Lα | F50CD2:Q70CG<br>3.17 | T98OG1:E55OE2<br>4.32 | 44.40 | 125.30 | 94.81 | 24.75c<br>19.78m | 1.25 | 32.62 | 1782 |
| 4H1Lβ | I49CD1:L60CB<br>3.30 | Y97CZ:D66CB<br>3.77 | 41.83 | 117.27 | 75.97 | 19.98c<br>18.84m | 1.06 | 27.16 | 1198 |
| 1ZGLα | K51NZ:D76OD1<br>6.91 | S91OG:E55OE2<br>2.90 | 51.82 | 117.60 | 85.36 | 26.11c<br>22.11m | 1.18 | 33.78 | 2831 |
| 1ZGLβ | N51CG:A64CB<br>3.70 | R99NH2:D66OD2<br>2.57 | 46.60 | 108.58 | 69.79 | 20.96c<br>14.06m | 1.49 | 25.67 | 1281 |

<sup>†</sup> Same TCR, peptide and DRA (DRB1-4) as 1fyt (DRB1\*0104).<sup>‡</sup> DRB1 numbering is +29 from conventional due to attached peptide; Q93=Q64; L96=L67; D95=D66; Q99=Q70; T106=T77.<sup>1</sup> Same as Table 1A legend220 221**Table 1C Summary Geometry TCR:pHLA-DR**

**Table 1C.** Summary Geometry TCR:pHLA-DR

| PDB | CDR2<sup>1</sup> | CDR3 | $\sigma^{\dagger}$ | $\lambda$ | $\omega$ | $\phi$ | $k^{\ddagger}$ | $\rho(\text{\AA})^{\diamond}$ | $dV^{\#}$ |
| --- | --- | --- | --- | --- | --- | --- | --- | --- | --- |
| 6CQL$\alpha^{\dagger}$ | L58CD1:A73CB<br>3.85 | A105N:E55OE1<br>3.00 | 37.14 | 133.17 | 93.59 | 20.41c<br>23.24m | 0.878 | 29.01 | 746 |
| 6CQLβ | S66OG:Q57NE2<br>3.01 | R108NH2:Y60OH<br>2.93 | 32.68 | 129.70 | 70.67 | 14.22c<br>18.95m | 0.750 | 25.56 | 542 |
| 6CQNα | L58CD2:T77CG2<br>3.73 | A105N:E55OE1<br>2.89 | 36.31 | 134.45 | 92.61 | 19.69c<br>22.34m | 0.881 | 28.98 | 690 |
| 6CQNβ | S66OG:Q57NE2<br>3.17 | L109CD1:D66CB<br>3.73 | 33.40 | 128.57 | 70.09 | 14.45c<br>23.56m | 0.613 | 25.68 | 586 |
| 6CQQα | L58CD2:A73CB<br>3.78 | A105N:E55OE1<br>2.88 | 35.95 | 134.32 | 89.29 | 18.85c<br>24.02m | 0.785 | 28.97 | 693 |
| 6CQQβ | S66OG:Q57OE1<br>3.64 | R108NH2:Y60OH<br>2.97 | 34.88 | 127.12 | 72.65 | 15.64c<br>22.76m | 0.687 | 26.34 | 800 |
| 6CQRα | T57OG1:Q70NE2<br>3.17 | A105N:E55OE1<br>3.03 | 36.52 | 133.75 | 87.44 | 18.75c<br>23.44m | 0.800 | 28.99 | 718 |
| 6CQRβ | E60OE2:K67NZ<br>3.07 | R108NH2:Y60OH<br>3.11 | 36.34 | 124.36 | 69.70 | 15.76c<br>29.07m | 0.542 | 25.95 | 746 |

<sup>†</sup> Same TCR (F24) and peptide and DRA (DRB1-11) as 6cqq (DRB1-15) and 6cqr (DRB1-1); same pMHC as 6cqn (TCR, F5; CDR3β G108R).<sup>1</sup> Same as Table 1A legend222

**Table 1D.**
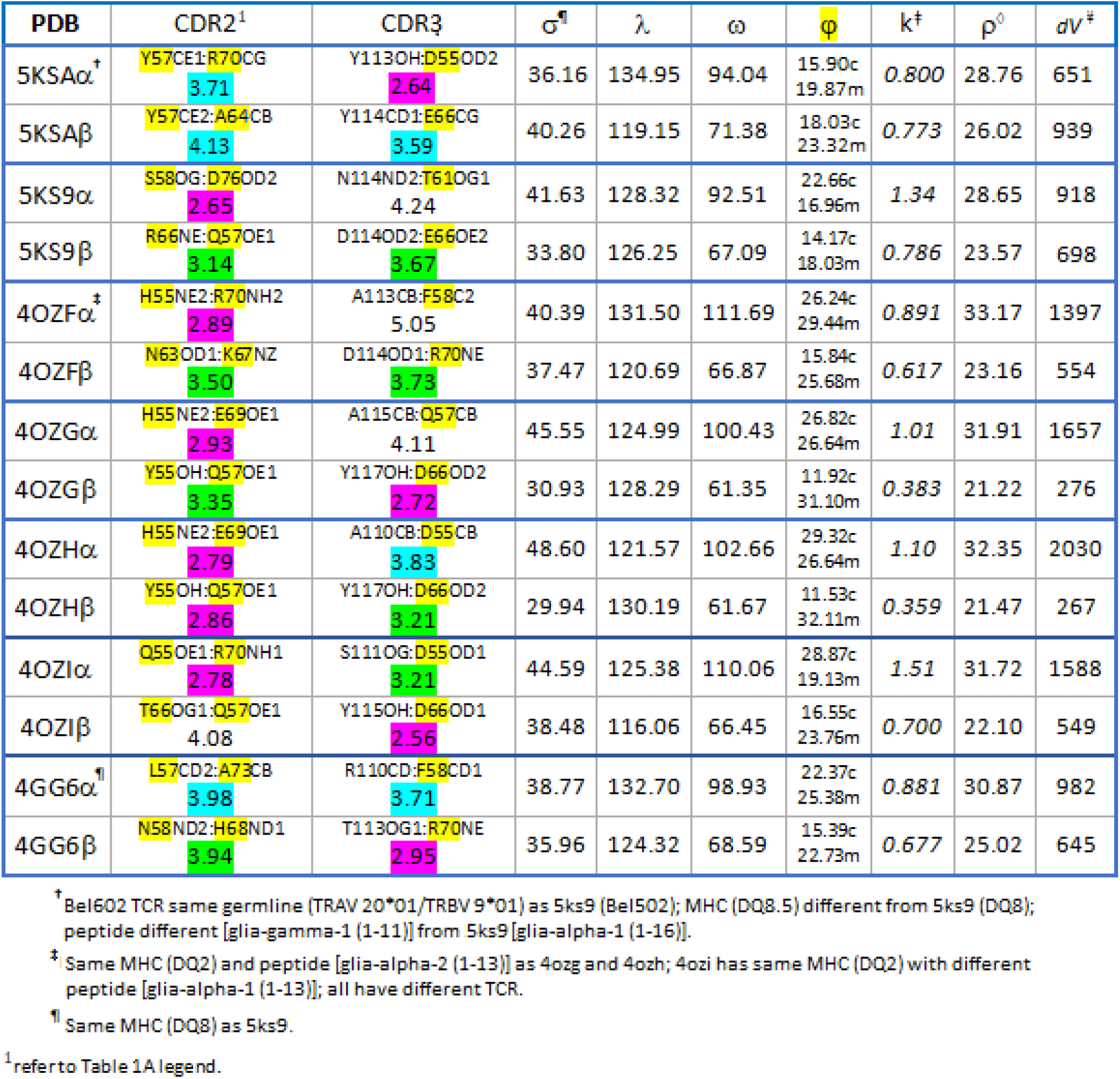
Summary of TCR:pHLA-DQ Geometry

Accordingly, the *mean dV* of these TCR V-domains was 955 Å^3^, excluding the unusually large *dV* of the 1ZGL Vα. Importantly, with the exclusion of 1ZGL (an apparent *outlier*) there is a linear relationship between calculated pitch (φ_c_) and the *dV* triple integral (see plotted values from Tables 1A⍰1D in the **Suppl.1.II**)—corresponding to the equation: *y = 83*.*60x* - *719*.*40;* where *R*^*2*^ *≈ 0*.*900*, by *linear regression* analysis. The lowest *dV* structures have the lowest calculated *pitch* values. Overall, it says something about TCR:pMHC that may not be intuitive—that a “flush” V-domain geometry could limit CDR2 “finding” an optimal binding interface with side-chains involving the aforementioned α-helix regions. This is the clonotypic nature of TCR function.

### CDR3:pMHC Contacts

All these structures, even those with the highest resolution, are subject to limitations of crystallography; and indeed, computed H-bonds are based on these coordinates. Nevertheless, a correlation between relative *dV* and contact distances for hydrogen bonds of the five components could be investigated; see Tables 2A⍰2C. For example, asking if particularly strong (close) H-bonds are linked to a restricted *dV*?

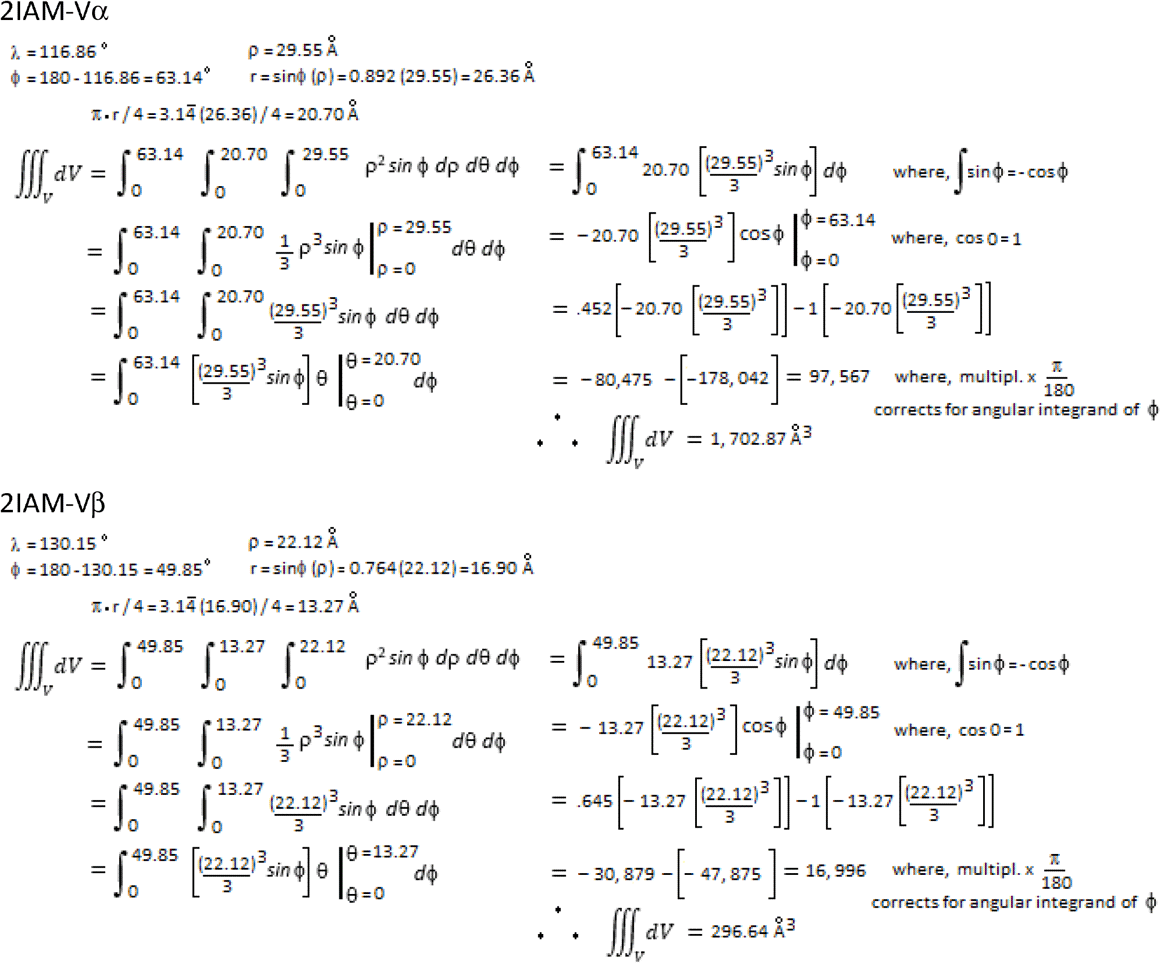

*for all (*n* = 38) triple-integral solutions, see **Suppl.1.I.**

**Table 2A.** Possible H-bonding networks in TCR:pHLA-DR complexes

| PDB | CDR3α:pep | Nn MHC:pep<sup>§</sup> | CDR3β:pep | pep:pep | $\alpha$:$\alpha$, $\alpha$:$\beta$, $\beta$:$\beta$ | MHC:MHC | $\alpha$/$\beta$:MHC |
| --- | --- | --- | --- | --- | --- | --- | --- |
| 2IAM* | none | A1N62ND2:L29O<br>2.0<br>A31N:A1N62OD1<br>2.2<br>A1N69ND2:A32O<br>2.1<br>V34N:A1N69OD1<br>1.8<br>B1R71NH1:N30O<br>2.1<br>B1R71NH2:N30O<br>2.0<br>N30ND2:B1Q70OE1<br>2.4 | T95OH:A31O<br>1.7<br>H96NE2:N30O<br>2.8<br>H96NE2:A31O<br>2.8 | none | αA93N:αQ91O<br>2.3<br>αQ94NE2:αA93O<br>2.1<br>αQ94NE2:αI90O<br>2.8 | B1R71NH1:<br>Q70OE1<br>2.2 | αQ91NE2:B1T77O<br>2.2<br>A1G58N:αG92O<br>2.4<br>βT98N:B1D66OD1<br>2.0 |
| 2IAN<br>(no H's) | none | A1N62ND2:L29O<br>3.0<br>A1N69ND2:A32O<br>3.1<br>V34N:A1N69OD1<br>3.0<br>B1R71NH1:N30O<br>2.9<br>B1R71NH2:N30O<br>2.8 | T95OH:A31O<br>2.5<br>H96NE2:N30O<br>3.8<br>H96NE2:A31O<br>3.8 | none | αG92N:αI90O<br>3.3 | B1R71NH1:<br>Q70OE1<br>3.2 | αQ91NE2:B1T77O<br>2.9<br>βT98OG1:D66OD1<br>2.7 |
| 4E41 | I28N:G91O<br>1.9 | A31N:A1N62OD1<br>1.7<br>A1N62ND2:L29O<br>2.0<br>A1N69ND2:A32O<br>2.1<br>V34N:A1N69OD1<br>2.0<br>B1R71NH1:N30OD1<br>2.1<br>B1R71NH2:N30O<br>1.8<br>B1Q70NE2:N30O<br>2.8 | R95NH1:I28O<br>1.9 | none | βR95NH2:αG91O<br>2.0<br>βR95NH2:αT93O<br>2.4<br>βR95NE:αD89OD1<br>2.5 | none | βT97N:D66OD2<br>1.9 |

<sup>§</sup> Nn, nearest neighbour; shown possible H-bonds, donor → acceptor; colour-code: $\leq 2.5$ , red; 2.6 to 2.9, mauve; $\geq 3.0$ , green.
\*Same TCR (E8) and MHC (DRA, DRB1\*0104; pep I28) as 2ian (pep T28); same pMHC as 4e41 (TCR, G4).

**Table 2B.**
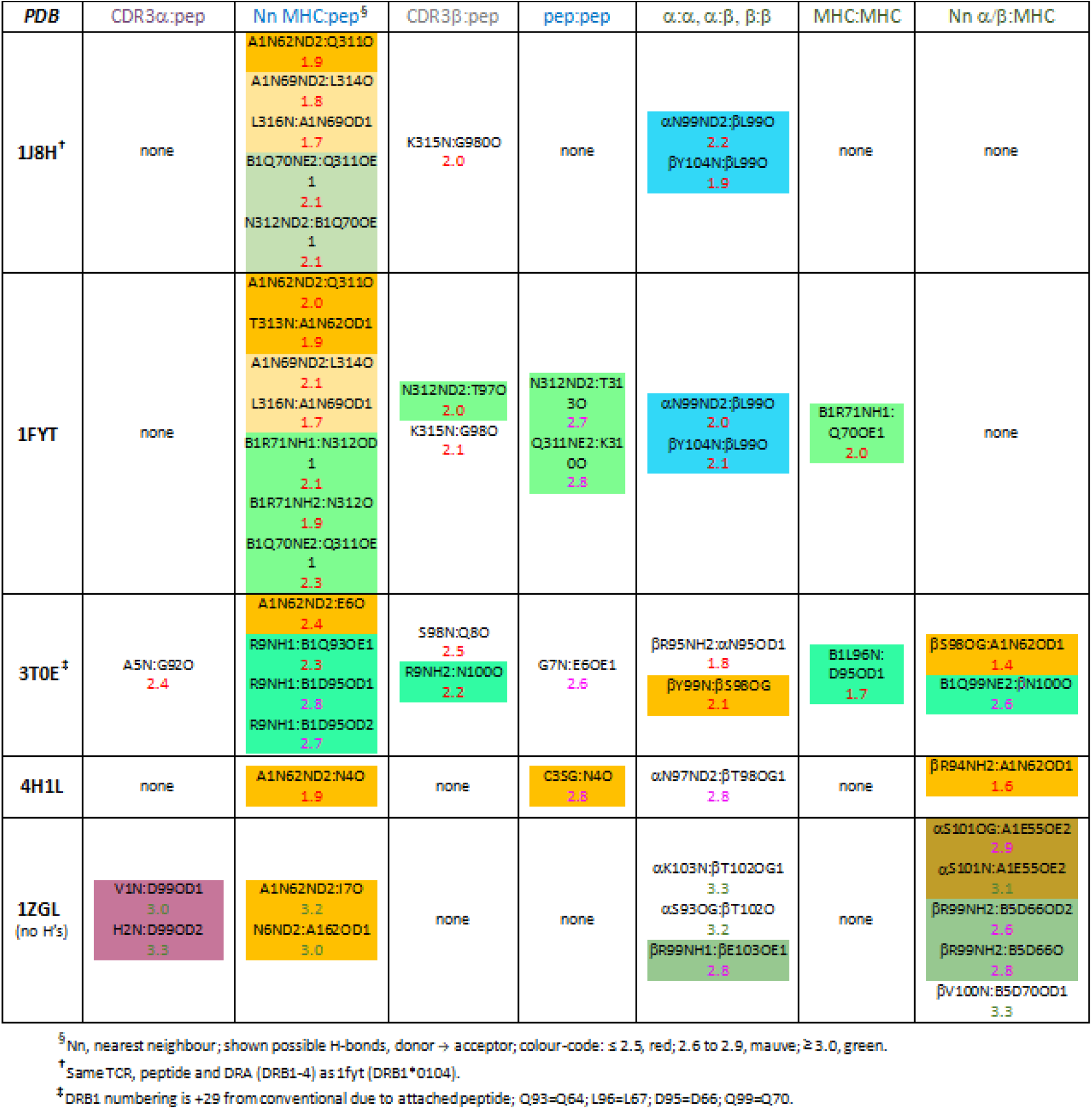
Possible H-bonding networks in TCR:pHLA-DR complexes

The structures were examined in *Swiss-PDB Viewer/Deepview-v4*.*1* (www.wpdbv.vital-it.ch); in *Swiss* H-bonds can be computed after hydrogens are added based on coordinates; two PDB files, (2IAN, 1ZGL) precluded adding hydrogens, and the shown H-bonding distances are thus ≈ 1 Å larger (Tables 2A, 2B). Here, an obvious feature of all the complexes is that there are two principal foci of H-bonding; one involving N62 of DRA (A1) (shaded *orange* in the tables), which may include a separate secondary grouping at N69 (*light-orange* shade), and the DRB1 (B1) centre involving R71 (*green* shades). Note, R71 is polymorphic, so structures involving allotypes/isotypes that have a different a.a. at pos. 71 (1J8H, 3T0E, 1ZGL, 6CQQ & 4H1L) involve a different a.a. (1J8H, 3T0E, 6CQQ), or simply do not have a corresponding beta-chain H-bonding centre (4H1L & 1ZGL). Superimposed upon the H-bonding centres are the H-bonds between the CDR3α and peptide, the CDR3β and peptide, any peptide to peptide H-bonding, any intra- or inter-bonds between the TCR V-domains, any MHC to MHC H-bonding, and direct H-bonds between TCR V-domains and the MHC (Tables 2A⍰2C, *columns*).

**Table 2C.** Possible H-bonding networks in TCR:pHLA-DR complexes

| PDB | CDR3α:pep | Nn MHC:pep<sup>§</sup> | CDR3β:pep | pep:pep | $\alpha\alpha$, $\alpha\beta$, $\beta\beta$ | MHC:MHC | Nn $\alpha/\beta$:MHC |
| --- | --- | --- | --- | --- | --- | --- | --- |
| 6CQL<sup>¶</sup> | R95NH1:A105O<br>2.0<br>R95NH1:G106O<br>2.6<br>R95NH1:N107OD1<br>2.3<br>R95NH2:N107OD1<br>2.0 | A1N69ND2:E97O<br>2.2<br>A99N:A1N69OD1<br>1.8<br>B1R71NH1:R95O<br>2.2<br>B1R71NH2:R95O<br>2.0 | A110N:E97OE1<br>1.9<br>G111N:E97OE2<br>1.9 | Q98N:E97OE1<br>2.0 | βG112N:βL109O<br>2.7 | B1R71NH2:<br>D70OD2<br>2.4 | αA105N:A1E55OE1<br>2.1 |
| 6CQN | R95NH1:A105O<br>2.1<br>R95NH1:G106O<br>2.6<br>R95NH1:N107OD1<br>2.2<br>R95NH2:N107OD1<br>1.9 | A96N:A1N62OD1<br>1.9<br>A1N62ND2:L94O<br>1.9<br>A1N69ND2:E97O<br>2.2<br>A99N:A1N69OD1<br>1.8<br>B1R71NH1:R95O<br>2.0<br>B1R71NH2:R95O<br>1.9 | A110N:E97OE1<br>1.7<br>G111N:E97OE2<br>2.0 | Q98N:E97OE1<br>2.2 | βG112N:βL109O<br>2.5 | B1R71NH2:<br>D70OD2<br>1.9 | αA105N:A1E55OE1<br>2.0 |
| 6CQQ | R95NH1:G106O<br>2.4<br>R95NH1:N107OD1<br>2.3<br>R95NH2:N107OD1<br>1.8 | A96N:A1N62OD1<br>1.8<br>A1N62ND2:L94O<br>1.8<br>A1N69ND2:E97O<br>2.1<br>A99N:A1N69OD1<br>2.0<br>S100N:B1D57OD1<br>2.6 | R95NH2:A110O<br>2.0<br>A110N:E97OE1<br>2.1<br>G111N:E97OE2<br>2.3 | Q98N:E97OE1<br>2.2 | βQ115NE2:αN114O<br>2.1 | none | αA105N:A1E55OE1<br>2.0<br>βR108NH2:B1D66OD1<br>2.2<br>βR108NH2:B1D66OD2<br>2.8 |
| 6CQR | R95NH1:A105O<br>2.2<br>R95NH1:N107OD1<br>2.1<br>R95NH2:N107OD1<br>2.0 | A96N:A1N62OD1<br>1.9<br>A1N62ND2:L94O<br>2.1<br>A1N69ND2:E97O<br>2.2<br>A99N:A1N69OD1<br>1.8<br>B1R71NH1:R95O<br>2.4<br>B1R71NH2:R95O<br>1.8 | R95NH2:A110O<br>2.2<br>A110N:E97OE1<br>1.8<br>G111N:E97OE2<br>2.2 | Q98N:E97OE1<br>2.4 | βQ115NE2:αN114O<br>2.1 | none | αA105N:A1E55OE1<br>2.0 |

<sup>§</sup> Nn, nearest neighbour; shown possible H-bonds, donor → acceptor; colour-code: $\leq 2.5$ , red; 2.6 to 2.9, mauve; $\geq 3.0$ , green.
<sup>¶</sup> Same TCR (F24) and peptide and DRA (DRB1-11) as 6cqq (DRB1-15) and 6cqr (DRB1-1); same pMHC as 6cqn (TCR, F5; CDR3β G108R).

In all cases, the CDR3α:peptide bonds are either non-existent, or isolated from the MHC:peptide centres. In 4E41, 6CQQ and 6CQR, CDR3α may link to CDR3β through both bonding nearby with peptide (*blue* shading). For 2IAM, 1J8H, and 1FYT there are bonds between intra-V (2IAM), or intra- and inter-V (1J8H, 1FYT), but these are not linked to the peptide (different *blue* shade). N62 is connected to a peptide:peptide H-bond (4H1L), or to an inter-Vβ bond (3T0E); in both cases these potential networks can include Vβ directly binding with N62 of DRA. 1ZGL is the only structure with an inter-V bond potentially linked to a direct V-domain interaction with MHC, while 4H1L is unique in not showing any DRB H-bonding involving the CDR3 (these two structures involve non-DRB1 isotypes). In 6CQL and 6CQN there is a potential DRB1:peptide link with an MHC:MHC bond, but this does not involve the TCR. The 6CQ series all have CDR3β:peptide bonds also involving a peptide:peptide bond; this is not found in any of the other complexes (*mauve* shading). Finally, and most significantly here, in 2IAM/2IAN, 1FYT and 3T0E the DRB1:peptide centre is potentially connected through a CDR3β:peptide bond with an MHC:MHC H-bond (Table 2A & 2B, *green* shades).

### CDR3:pMHC Chemistry

H-bonds computed in *Swiss* are potential H-bonds, and to assess a given network the chemistry of each bond must be examined [24, 25]. The analyses of Tables 2A⍰2C indicated that a potential H-bonding network where CDR3β is connected through peptide-bonding to DRB1:peptide and DRB1:DRB1 H-bonds might correlate with the *highly-restricted dV* of Vβ in 2IAM (297 Å^3^) and 2IAN (270 Å^3^). However, these networks would need to be distinct from this same type of potential network observed in 1FYT and 3T0E (Table 2B), which have *dV* for Vβ of 1747 Å^3^ and 1024 Å^3^, respectively (Table 1B). Shown is the analysis of the 2IAM-Vβ versus 1FYT-Vβ for contacts made by each CDR3β (Fig. 3). For 2IAM-Vβ a potential H-bonding network involving CDR3β a.a. Y95 and H96 focused on central peptide a.a. N30 and A31 is apparent; and DRB1 a.a. Q70 and R71 appear in this same network. More specifically, there are three *potential* H-bonds with the N30:O (at R71:NH1, R71:NH2, and H96:NE2); Y95 appears to H-bond to A31:O, and there is also a potential bond between Q70:OE1 and R71:NH1 (Figs. 3A & D). By contrast, 1FYT-Vβ displays single H-bonds between T97:O and N312:ND2 (peptide), and G98:O and K315:N (peptide). The DRB1, R71:NH2 is involved with H-bonding to N312:O, and R71:NH1 has possible H-bonds with N312:OD1 and with Q70:OE1 (Figs. 3B & E). Since 1FYT and 1J8H only differ via DRB1 alleles, we compared H-bonding for the two (Fig. 3B *vs*. 3C; 3E *vs*. 3F). Illustrating the importance of the *Swiss* computation, in Figure 3C it was anticipated that the R71K polymorphism would effect three H-bonds—involving K71 in a manner similar to R71. However, as is shown in Figure 3F, R71K actually shifted peptide contacts to Q70; in-fact, K71 did not show any H-bonds. Importantly, this confirms the change in peptide conformation between the two structures as originally reported [13], and suggests that alloreactivity of 1FYT/1J8H involves the single G98:0 to K315:N bond in maintaining the ≈ 1740 Å^3^ *d*V of the two Vβ (Table 1B). Still, the chemistry of each possible bond must be examined to arrive at feasible mechanisms; arguably, this is necessary to understand binding.

**Figure 3.**
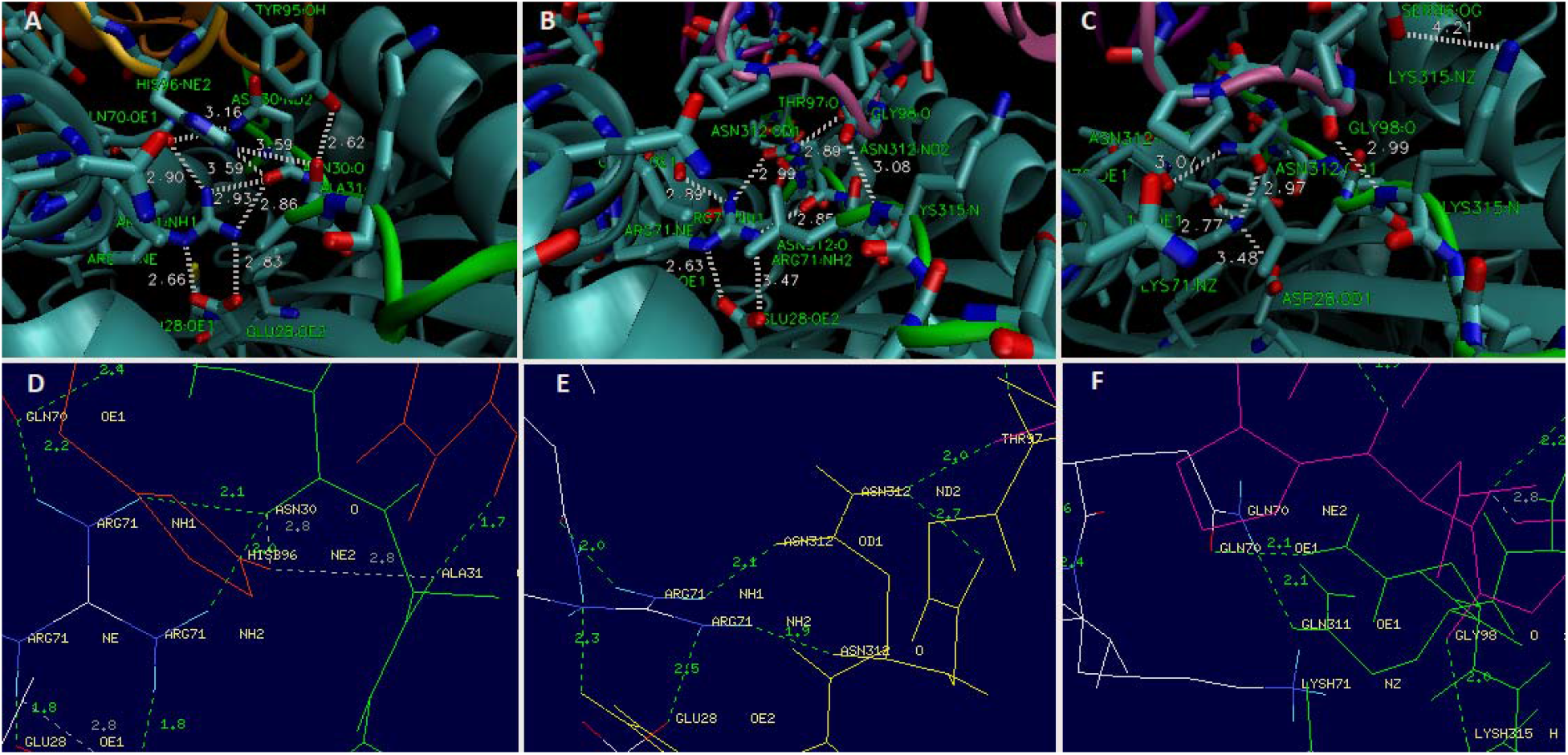
CDR3 H-bonding networks of 2IAM-Vβ (A & D), 1FYT-Vβ (B & E) and 1J8H-Vβ (C & F). *PDB* files analysed with *VMD* (top) and *Swiss-Deepview* (bottom). In *Swiss*, hydrogens added and H-bond distances computed; see *dotted* lines in *green*. Involved side chains shown by atom in *licorice* (top) or *stick* (bottom); The 2IAM CDR3β in *gold* (A) and *red* (D); 1FYT/1J8H CDR3β in *mauve* (B, C, E, F); and peptides in *green*. Figure (A⍰C) are the original output of the indicated *PDB* file as analysed in *VMD-1*.*9*.*1*. Figure (D⍰F) are the original *PDB* outputs as analysed in *Swiss-Deepview-v4*.*1*. Best viewed at 125%.

### H-Bonding Mechanisms

Shown in Figure 4A is a suitable reaction mechanism for the 2IAM CDR3β binding to the peptide (trios phosphate isomerse, 15mer)-HLA-DR complex (*PDB* ref., 15). When considering the probability of hydrogen bonds in a mechanism, all possibilities were drawn using *ChemSketch* (ACD Labs; www.acdlabs.com/resources) and electron pathways traced using standard evaluations of electron configuration [24, 25]. Thus DRB1 (B1) Q70:OE1 would be in an H-bond with B1R71:NH1 in the absence of the peptide, with the downstream effect of relieving the charge on the B1R71:NH2. Bound peptide has the same effect on R71:NH2, where the N30:O attacks the R71:NH2 (*orange* arrows). Here, the peptide would indirectly break the H-bond between Q70 and R71 (*orange* blocks). When the TCR binds, the βH96:NE2 charge is preferentially attacked by the N30:O (*purple* arrows); this reverses the previous bond between the peptide and DRB1 (*purple* block) and has the downstream effect of re-forming the intra-MHC bond between Q70 and R71 (*purple* blocks on the *orange* blocks). Note, this is the only obvious mechanism that relieves both the H96 and R71 charges, and would be favoured over an A31:O attack on H96:NE2 by the neighbouring βY95 H-bond with A31:O (Fig. 4A). Therefore, *just three* of the possible H-bonds are predicted; and the overall effect is a charge-relay [28, 32, 33] between the MHC, the peptide, and the TCR.

**Figure 4.**
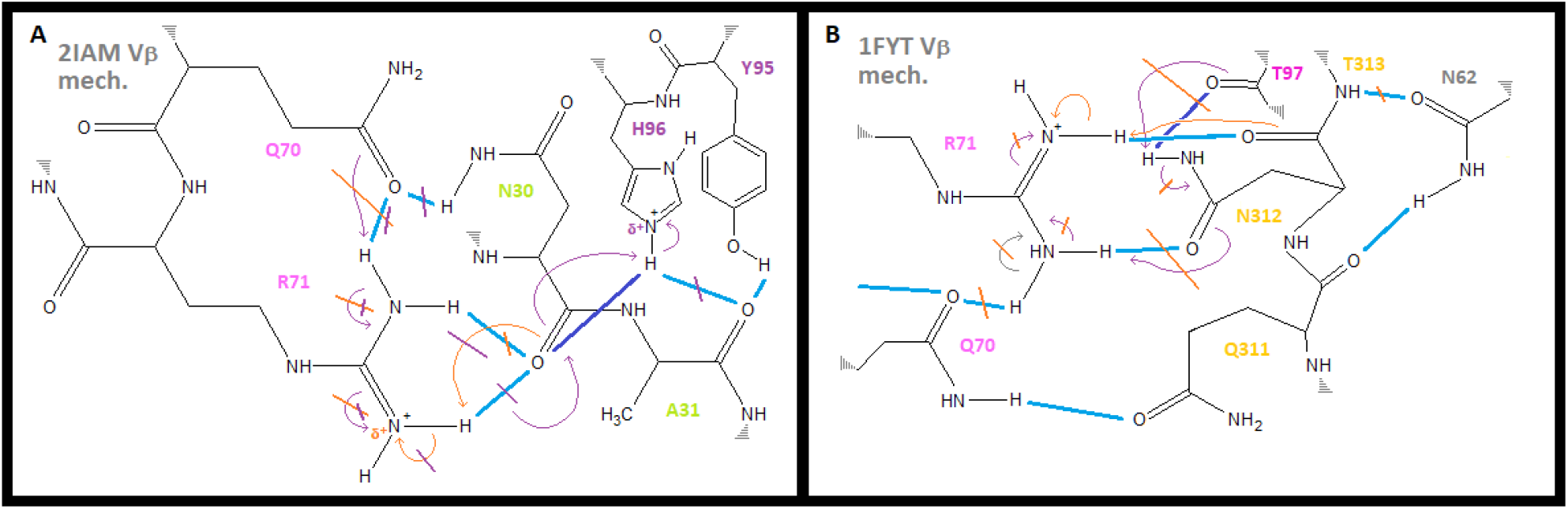
H-bonding mechanism for CDR3β binding to peptide and MHC. *ChemSketch* drawn mechanisms are shown for 2IAM (A) and 1FYT (B) and were based on analyses in *VMD* and *Swiss* (Fig. 3). *Not necessarily to scale*. Electrons shown with arrows (*purple*, involving the TCR; *orange*, related to the peptide). Possible H-bonds are shown in *blue* (darker shade for key TCR bond) and are crossed with the color-coded *block* line if not probable (see text). Best viewed at 125%.

Physiologically, the charge on R71:NH2 could be initially neutralized via the neighbour Q70:OE1—subsequently replaced during processing by CLIP, and/or at HLA-DM exchange, with antigenic peptide (ref. 9). During T-cell conjugation with the antigen presenting cell, the TCR would replace the A30:O to R71:NH2 attack with an A30:O to H96:NE2 attack, which *relays* the charge back to R71:NH2—subsequently, neutralized again by Q70:OE1. While the hallmark charge-relay network of the ‘catalytic triad’ within serine proteases stabilizes formation of a covalent acyl-enzyme intermediate [24]; here, a relay is utilized in a similar (albeit, noncovalent) role. Undoubtedly, this would still be a particularly stabilised transition-state, *i*.*e*., given that these H-bonds are de-localised across 3/5 components of the structure [24⍰27].

By contrast, Figure 4B shows a suitable mechanism for the analogous CDR3β binding reaction with the influenza HA peptide-HLA-DR complex of 1FYT. The differences are the peptide and TCR, as both 2IAM and 1FYT are DRB1*0104 structures. Similarly the peptide would replace the internal B1Q70:OE1 to B1R71:NH1 with an H-bond directly with the charged R71:NH2 via the peptide N312:O (*orange* arrows). Again, the TCR could replace this bond with an attack by the CDR3β (here, T97:O on N312:ND2; *purple* arrows). However, the downstream effect of such an attack in this case is a partial charge on N312:OD1 and an attack on R71:NH1. Note this would restore the charge on R71:NH2, effectively breaking the peptide-MHC bond. Also, despite what looks like proximity in the *ChemSketch* diagram, T97 is actually far removed from R71 (Fig. 3E) such that an attack directly on R71:NH2 is not possible. Thus, because of a remaining charge on R71:NH2, the T97:O attack on N312:ND2 (with its downstream effects) would *not* be favoured (*orange* blocks). While similarly to 2IAM there is a neighbouring H-bond (in this case to Q311), the net loss of a CDR3β H-bond clearly distinguishes the two mechanisms, and shows that the 1FYT CDR3β does not favour a charge-relay mechanism; *i*.*e*., would not have the stabilized CDR3β tether at B1R71—ostensibly, essential chemistry driving restricted *dV* in 2IAM. Note that the 2IAN-Vβ mechanism appears to be identical to that of 2IAM (structures differ by a single, sufficiently distant, a.a. of the peptide). Next, 3T0E also showed the TCR:peptide:MHC:MHC type of potential network by contacts analysis (Table 2B). Figure 5A shows the a.a. involved in this potential network using *Swiss*, and the proposed mechanism is shown in Figure 5B. Note that the DRB1-4 allotype of 3T0E has the R71K polymorphism, and that the involved peptide a.a. is R9. In terms of distinguishing the mechanism from 2IAM, the question is whether the CDR3β N100 H-bond to R9 could exist with a B1:peptide H-bond. In fact, if the Vβ closest contact N100:O is to attack the positive charge of R9:NH2 (purple arrows), it would have to compete (indirectly) with the H-bond of B1Q93:OE1 to R9:NH1 (orange arrows). Together with the B1Q99:NE2 possible bond to the same N100:O, the B1Q93 bond would be favoured; thus, the 3T0E mechanism is not similar to 2IAM, most notably because CDR3β is not likely to form an H-bond with the peptide. Note that in the related structure, *PDB* 3O6F, this conclusion holds as well, where B1D95:OD2 has a possible H-bond to R9:NH2 at 1.6 Å; again, not favouring a CDR3β N100:O bond to the peptide [28].

**Figure 5.**
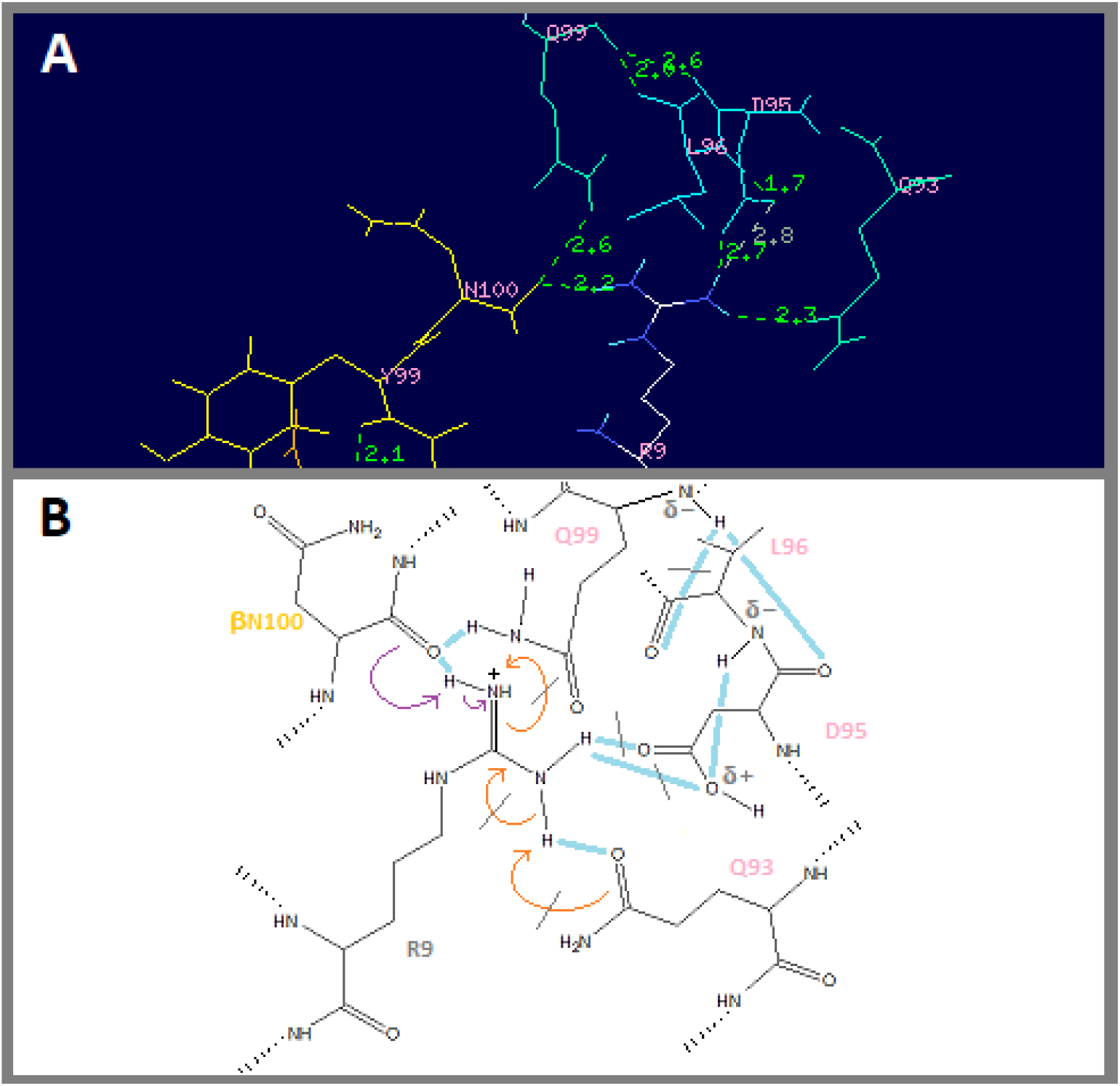
H-bonds and H-bonding mechanism for 3T0E CDR3β. *Swiss* was used as before (non-involved a.a. deselected for simplicity) to compute possible H-bonds for 3T0E (A). Mechanism for 3T0E TCR binding via CDR3β to peptide and DRB1 (B) shows possible H-bonds in *blue*, with improbable bonds blocked with *grey* lines (*ChemSketch*). Electron flow shown with *purple* arrows (TCR), or orange (MHC). CDR3β in *yellow* (top) or a.a. labelled *yellow* (bottom); DRB1 in *mauve* (top) or labelled a.a. in *pink* (bottom); peptide by atom (top) or a.a. labelled *grey* (bottom). Figure (A) is the original output of the *PDB* file as analysed in *Swiss-Deepview-v4*.*1*. Figure (B) is a diagram constructed in *ChemSketch*. Best viewed at 100%.

### DQ H-Bonding Mechanisms

As shown in Table 1D, we noted the highly-restricted *dV* in Vβ for two of the TCR:pHLA-DQ structures (4OZH & 4OZG; 267 Å^3^ and 276 Å^3^, respectively) [*PDB* ref. 29]. Here, the control structure is 4OZF (Vβ *dV*, 554 Å^3^), where the pMHC is the same; in-fact, these three structures only differ by their CDR3 regions (*all 38 IMGT-computed CDR3 junctions shown in Suppl*.*1*.*III*). Accordingly, H-bonding was analysed with *Swiss-Deepview* as before (Fig. 6). While very similar, we noted the absence (Fig. 6A & B) of the DQA1, T61:OG1 role in H-bonding to Vβ, R109:NH1 (Fig. 6C). This is accomplished in 4OZH by preferential H-bonding to Vβ a.a. S110 (Fig. 6A), or sterically in 4OZG by Vβ a.a. F110 (Fig. 6B). Thus, only in 4OZF is T61:OG1 free to H-bond to the pivotal arginine (CDR3β, R109); note, this shift from R109 contacting MHC at a.a. N62 to T61 indirectly disrupts the MHC:MHC component of the charge-relay mechanism (Fig. 7C *vs*. 7A & B). Crucially, the chemistry of S110 versus F110 versus A110 in these TCR is ultimately the result of somatic genetic construction of the CDR3β loop [**Suppl.1.III**; ref. 4].

**Figure 6.**
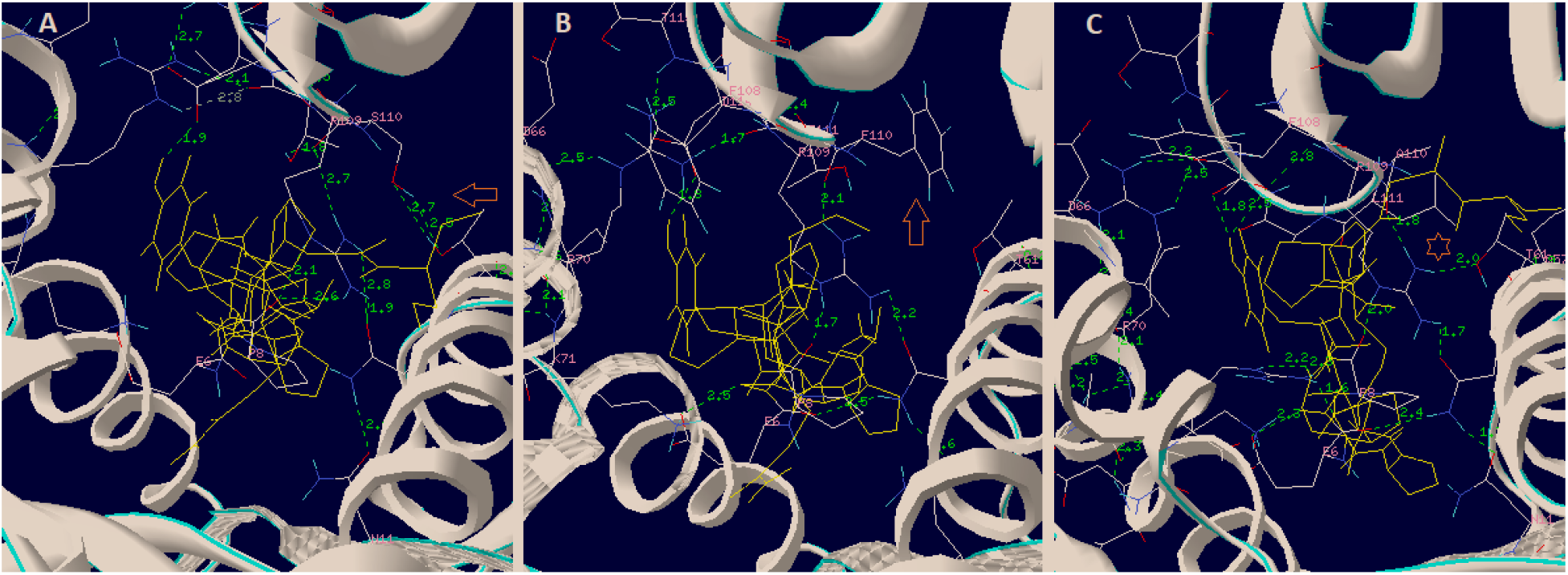
H-bonds for three different TCR CDR3β, (from germline-identical TCR) on the same pHLA-DQ. (A) 4OZH, showing S110:T61 H-bonds (*orange* arrow), (B) 4OZG, showing bulky F110 (orange arrow); both of which could preclude (C) 4OZF, the conserved R109 H-bonds to T61 (*orange* star). *Swiss-Deepview* analysis as previously; DQ and CDR3β backbones in *white* ribbon; peptides in *yellow* wire; involved side chains in *CPK*; computed possible H-bonds and distances in *green*. Figures are the original output of the indicated *PDB* files as analysed in *Swiss-Deepview-v4*.*1*. Best viewed at 150%.

**Figure 7.**
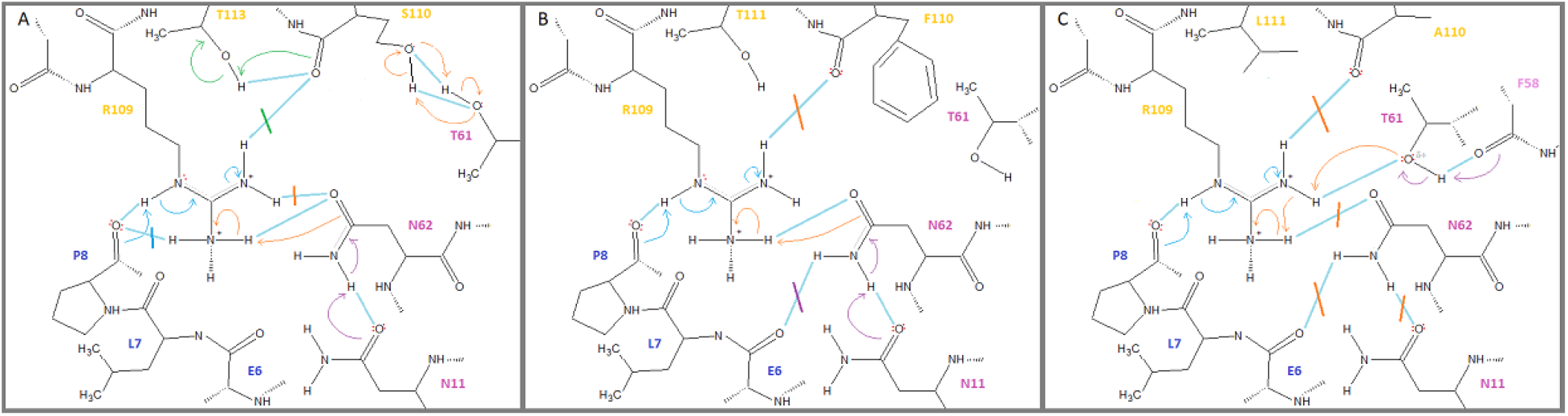
H-bond mechanisms for three different CDR3β, (from germline-identical TCR) on the same pHLA-DQ. (A) 4OZH, showing S110:T61 H-bonds (*orange* arrows), (B) 4OZG, showing bulky F110; both of which could preclude (C), the conserved R109 H-bonding to T61, as only observed for the nominal *d*V Vβ of 4OZF. As previously, possible H-bonds in *blue*, with improbable bonds blocked with *colour-matched* lines (*ChemSketch*). Electron flow shown with *blue* arrows (peptide:TCR), or *orange* arrows (MHC:TCR), or *purple* arrows (MHC:MHC). CDR3β a.a. labels in *yellow*, peptide a.a. in *blue*, DRB1 a.a. in *mauve*. Best viewed at 150%.

A sequentially mechanism can be envisioned. First, there is internal MHC:MHC H-bonding between N11 and N62 (*purple* arrows, Fig. 7A & B). This facilitates the TCR binding to peptide via a P8:O attack on the R109:NE proton (*blue* arrows), and this leads to neutralizing the charge on R109:NH2. Uniquely here (*purple* arrows, Fig. 7C), when the CDR3β is permissible for the T61:OG1 to R109:NH1 H-bond, *i*.*e*., the loop a.a., A110-L111-A112-A113 (4OZF), this appears to preclude both N62:OD1 attack on the charged R109:NH2 and the MHC-MHC H-bond (N11:OD1 to N62:ND1); compare in Fig. 7C to 7A & B). Thus, all three TCR (across DR and DQ) which displayed highly-restricted *dV*, also shared the likelihood of a charge-relay mechanism of CDR3β binding, where an MHC to MHC bond is involved.

## Discussion

We used spherical coordinates to derive a unique solved triple integral for each of the 38 V-domains (Figs. 1 & 2; **Suppl.1**.*I*). Using this approach the mean volumetric-density of a TCR V-domain through the putative CDR2 scanning path was 1060 Å^3^. As indicated, ⍰⍰⍰ *dV* is a ‘slice’ volume element of a cone with the vertex at the ‘across-the-groove’ MHC α-helix, where maximal CDR2 scanning (*d*θ) calculates close to the actual range of α-helix distances between the most N-term. and most C-term. CDR2 contacts (Tables 1A⍰1D). Also, overall geometry consistent with MHC tethering via CDR3 (first formulated for TCR-Vα:pHLA-A2; ref. 20) was again, broadly apparent. Most importantly, here a linear relationship was found between V-domain *pitch*; calculated by:

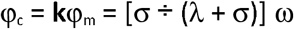

and V-domain “*dV*” calculated by:

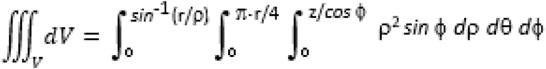

This suggests that reduced pitch limits *d*θ, *i*.*e*., the CDR2-scanning function. In retrospect, this might confirm intuition (excluding the broader *TCR-CD3 complex*, of course) on possibilities for a mechanism involving just the five components. Indeed, H-bonding chemistry which effects relatively simple physics is a hallmark of quite diverse protein machinery [24, 30]. Also, TCR had *k* values that varied in either direction—indicating that conformation adjustments might moderate flush and open pitch *without* much increasing or decreasing of the *dV, viz*., *R*^2^ ≈ *0*.*900* (**Suppl.1.II**). Thus, pitch calculated from a given V-domain’s *twist*-*tilt*-*sway* (orientation) might be a kind of “hidden” correlate of TCR-selection; although there is not available crystallography on any thymic (selecting) ligands [31]. For 2IAM, there are data for the uncomplexed TCR, and the CDR3β backbone is displaced ≈ 3.4 Å upon binding pMHC, while βY95 moves ≈ 9.0 Å to form the H-bond shown (Fig. 3A); indeed, the key to understanding the TCR may be in this relationship between two induced-fit TCR conformations [32⍰35]. Suggested here is a conservation/approximation of *dV* for V-domains in TCR binding the *selecting* (thymus) and *activating* (peripheral) pMHC. As shown (Fig. 3E *vs*. 3F), alloreactivity displays shared *dV* accomplished by quite different CDR3 binding chemistry. Alternatively/historically, *intrinsic* TCR affinity (*i*.*e*., equilibrium and rate binding constants specified by the α/β TCR protein) has been used to explain both thymic TCR selection and peripheral TCR recognition [4, 36, 37]. Clearly TCR distinguish rare target (agonist) pMHC from thousands of nonagonist pMHC on the APC surface; and this is not mutually exclusive of whether or not a physical force (TCR loading) is an obligate component of TCR function [8, 37]. In these and other regards, there is a stereochemical alternative to an affinity-limited binding reaction [25⍰27].

Briefly, we assume TCR:pMHC reactions involve a high(er) energy “scanning” conformer— because, scanning (leading to a suitable CDR2:MHC interface) has the effect of lowering the transition-state free energy, ΔΔ*G*^⍰^ (*i*.*e*., *Curtin*-*Hammett* control) [25, 27, 38]. For example, in TCR like E8 (of 2IAM/2IAN) very little CDR2β scanning is apparently needed, due to the effect of the proposed charge-relay stabilised CDR3β:pHLA-DR transition-state. Thus, *dV* is a universal consequence of CDR3 chemistry; but usually the reaction requires more CDR2-scanning (*i*.*e*., usually V-domains have larger *dV* values; Tables 1A⍰1D). Recently, ensemble refinement of crystal data with MD simulations has suggested conformational diversity in the microsecond range, but the conformational changes implicated here-in would be on the millisecond to minutes scale; in this regard, NMR of membrane-bound receptors offers promise [39], and may elucidate the potential of internal water(s) in these H-bonding networks [32]. Finally, spherical coordinates have been used by others to calculate TCR *center-of-mass* variation in TCR:pMHC complexes [40]. These investigators did not consider Vα and Vβ independently, although *Hoffmann et al*., had shown the angle between Vα and Vβ characterized different TCR in pMHC complexes; allowing these researchers to group a panel of TCR into six different clusters linked to clonotype [41]. While there are myriad downstream implications, taken together, these data support that V-domain rotation and germline to germline contacts between TCR and pMHC *both* depend upon CDR3 H-bonding with highly-conserved MHC α-helix motifs. Thus, unlike TCR-affinity, V-domain *dV* is clonotypic. In this regard, similar to the class-I, R65-motif [20, 42], the class-II motifs presented here are found in *Galago sp*. (NCBI Acc. No. AAA96291) as well as in both *tarsiers* and *lemurs*; suggesting conservation for at least 63 million years [43, 44].

## Methods

### Availability of data and materials

*PDB* files are public and available at www.ncbi.nlm.nih.gov; all analytics data are either in the paper, or supplementary materials; *Competing interests*: Dr. Murray and *Xenolaüs Genetics LLC* declare no competing interests regarding the research, or its publication; *Author Contributions*: J.S.M. did the research and wrote the manuscript.

### *PDB* Analysis

*VMD-1*.*9*.*1* software (www.ks.uiuc.edu) was used for *PDB* files downloaded via *NCBI* (www.ncbi.nlm.nih.gov) from the *RSCB-PDB* (www.rscb.org); views normalized with the *VMD xyz-axis* tool; alpha carbon (Cα) main-chains in *new cartoon*; all alleles named per *NCBI* annotation. *Euler*’s methods (www.mathword.wolfram.com/EulerAngles.html) were the basis for the specific angle analyses, as previously reported. Briefly, three angles corresponding to the *twist, tilt*, and *sway* of each domain over the pMHC were measured from fixed Cα through the 19 structures: (i) in-plane to the MHC-groove {*twist* = ω}, with (ii) displacements perpendicular to the groove {*tilt* = λ}, including (iii) side-to-side variation {*sway* = σ}. The a.a. positions used as coordinates for angular measures across structures were *fixed*; see previous. The incline of a V-domain, {*pitch* = φ_m_}, was approximated (calculated) by the equation: φ_c_ = [σ ÷ (*λ* + σ] ω (see also **Results**, Tables 1A⍰1D). Pitch was also measured by using the closest determined CDR2 contact Cα for an angle across-the-groove to the closest CDR3 contact with an α-helix side-chain (vertex), then back to said CDR2 closest contact within the opposite α-helix (≈ 2-fold symmetry); angular value in degrees via the *VMD angle-label* tool. Triple integrals for all 38 V-domains based on the fixed geometry were solved as described in **Results** (**Suppl.1.I**). Linear regression analysis by *MS-Excel* (**Suppl.1.II**); and statistics by paired two-tailed *Student*’s t-test (www.insilico-net/tools/statistics/ttest).

### Contacts Analysis

All measures were performed with *Swiss* or *VMD-1*.*9*.*1* as is specified in *Results*. Closest contacts in *angstroms* (Å) were determined by examining appropriate coordinates between structures (computed in *Swiss*, or measured/computed in *VMD*). Individual atomic contacts are named per software annotation. DRA chain contacts are *abbrv*. “A1” to avoid confusion with single-letter a.a. code for alanine (A); potential H-bond networks are colour-coded as is described in table captions and the text.

### CDR3 Joint Analysis

Nucleotide sequences for all CDR3 of TCR were specified from *PDB* files (*Suppl*.*1*.*III*). TCR a.a. sequences were reverse translated using the *SMS* tool at www.bioinformatics.org. These were then imported into *IMGT* algorithms for joint analysis (www.imgt.org). Amino acid sequences of resulting CDR3 joints were determined by the *IMGT* algorithm; consensus *IMGT* numbering.

### H-Bonding Mechanisms

Hydrogen (H) bonds were estimated either with *VMD*, or (more accurately) with *Swis*s. In *Swiss*, H-bonds are computed after computing hydrogens to the structures. H-bond distances were used as a factor in determining suitable organic reaction mechanisms, where relevant side-chains were reproduced with *Chemsketch* (www.acdlabs.com/resources). Standard evaluations of electron configuration per relevant atoms were used to predict electron flows [25].

## Acknowledgements

Crystallographers, molecular biologists and computer scientists providing public availability of coordinates, sequences and software made this work possible. Thank you.

## Supplementary details

**[Suppl.1.pdf]**

I. Triple Integral Solutions for all V-domains

II. Linear Regression Analysis of Calculated Pitch versus *dV* (all V-domains)

III. *IMTG* CDR3 Joint Analysis (all V-domains)

