## Supplement 1 for "CDR3 binding chemistry controls TCR V-domain rotational probability and germline CDR2 scanning of polymorphic MHC"

### Supplementary details for:

### Somatic genetics of CDR3 control TCR V-domain rotational probability effecting germline CDR2 scanning of polymorphic MHC

Title: **Supplement 1**

#### I. Triple Integral Solutions for V-domain Panel

#### II. Linear Regression Analysis of Calculated Pitch versus *dV* (V-domain Panel)

#### III. *IMTG* CDR3 Joint Analysis (V-domain Panel)

Author: **Joseph S. Murray, Ph.D.**

#### Supplement 1.I. Triple Integral Solutions for all V-domains

$\lambda = 116.86°$ $\rho = 29.55\ Å$

$\phi = 180 - 116.86 = 63.14°$ $r = \sin\phi\ (\rho) = 0.892\ (29.55) = 26.36\ Å$

$\pi \cdot r / 4 = 3.1\bar{4}\ (26.36)/4 = 20.70\ Å$

**2IAM-Vα**

$$\iiint_V dV = \int_0^{63.14}\int_0^{20.70}\int_0^{29.55} \rho^2 \sin\phi\ d\rho\ d\theta\ d\phi \quad = \int_0^{63.14} 20.70\left[\frac{(29.55)^3}{3}\sin\phi\right] d\phi \quad \text{where, } \int \sin\phi = -\cos\phi$$

$$= \int_0^{63.14}\int_0^{20.70} \frac{1}{3}\rho^3 \sin\phi \Big|_{\rho=0}^{\rho=29.55} d\theta\ d\phi \quad = -20.70\left[\frac{(29.55)^3}{3}\right]\cos\phi \Big|_{\phi=0}^{\phi=63.14} \quad \text{where, } \cos 0 = 1$$

$$= \int_0^{63.14}\int_0^{20.70} \frac{(29.55)^3}{3}\sin\phi\ d\theta\ d\phi \quad = .452\left[-20.70\left[\frac{(29.55)^3}{3}\right]\right] - 1\left[-20.70\left[\frac{(29.55)^3}{3}\right]\right]$$

$$= \int_0^{63.14}\left[\frac{(29.55)^3}{3}\sin\phi\right]\theta \Big|_{\theta=0}^{\theta=20.70} d\phi \quad = -80{,}475 - \left[-178{,}042\right] = 97{,}567 \quad \text{where, multipl. x } \frac{\pi}{180} \text{ corrects for angular integrand of } \phi$$

$$\therefore \iiint_V dV = 1{,}702.87\ Å^3$$

$\lambda = 130.15°$ $\rho = 22.12\ Å$

$\phi = 180 - 130.15 = 49.85°$ $r = \sin\phi\ (\rho) = 0.764 (22.12) = 16.91\ Å$

$\pi \cdot r / 4 = 3.1\bar{4}\ (16.90)/4 = 13.28\ Å$

**2IAM-Vβ**

$$\iiint_V dV = \int_0^{49.85}\int_0^{13.28}\int_0^{22.12} \rho^2 \sin\phi\ d\rho\ d\theta\ d\phi \quad = \int_0^{49.85} 13.28\left[\frac{(22.12)^3}{3}\sin\phi\right] d\phi \quad \text{where, } \int \sin\phi = -\cos\phi$$

$$= \int_0^{49.85}\int_0^{13.28} \frac{1}{3}\rho^3 \sin\phi \Big|_{\rho=0}^{\rho=22.12} d\theta\ d\phi \quad = -13.28\left[\frac{(22.12)^3}{3}\right]\cos\phi \Big|_{\phi=0}^{\phi=49.85} \quad \text{where, } \cos 0 = 1$$

$$= \int_0^{49.85}\int_0^{13.28} \frac{(22.12)^3}{3}\sin\phi\ d\theta\ d\phi \quad = .645\left[-13.28\left[\frac{(22.12)^3}{3}\right]\right] - 1\left[-13.28\left[\frac{(22.12)^3}{3}\right]\right]$$

$$= \int_0^{49.85}\left[\frac{(22.12)^3}{3}\sin\phi\right]\theta \Big|_{\theta=0}^{\theta=13.28} d\phi \quad = -30{,}902 - \left[-47{,}911\right] = 17{,}009 \quad \text{where, multipl. x } \frac{\pi}{180} \text{ corrects for angular integrand of } \phi$$

$$\therefore \iiint_V dV = 296.86\ Å^3$$

$\lambda = 134.93°$ $\rho = 29.77\ Å$

$\phi = 180 - 134.93 = 45.07°$ $r = \sin\phi\ (\rho) = 0.701 (29.77) = 20.87\ Å$

$\pi \cdot r / 4 = 3.1\bar{4}\ (20.87)/4 = 16.39\ Å$

**1J8H-Vα**

$$\iiint_V dV = \int_0^{45.07}\int_0^{16.39}\int_0^{29.77} \rho^2 \sin\phi\ d\rho\ d\theta\ d\phi \quad = \int_0^{45.07} 16.39\left[\frac{(29.77)^3}{3}\sin\phi\right] d\phi \quad \text{where, } \int \sin\phi = -\cos\phi$$

$$= \int_0^{45.07}\int_0^{16.39} \frac{1}{3}\rho^3 \sin\phi \Big|_{\rho=0}^{\rho=29.77} d\theta\ d\phi \quad = -16.39\left[\frac{(29.77)^3}{3}\right]\cos\phi \Big|_{\phi=0}^{\phi=45.07} \quad \text{where, } \cos 0 = 1$$

$$= \int_0^{45.07}\int_0^{16.39} \frac{(29.77)^3}{3}\sin\phi\ d\theta\ d\phi \quad = .706\left[-16.39\left[\frac{(29.77)^3}{3}\right]\right] - 1\left[-16.39\left[\frac{(29.77)^3}{3}\right]\right]$$

$$= \int_0^{45.07}\left[\frac{(29.77)^3}{3}\sin\phi\right]\theta \Big|_{\theta=0}^{\theta=16.39} d\phi \quad = -101{,}765 - \left[-144{,}143\right] = 42{,}378 \quad \text{where, multipl. x } \frac{\pi}{180} \text{ corrects for angular integrand of } \phi$$

$$\therefore \iiint_V dV = 739.64\ Å^3$$

$\lambda = 109.62°$ $\rho = 27.91\ Å$

$\phi = 180 - 109.62 = 70.38°$ $r = \sin\phi\ (\rho) = 0.942 (27.91) = 26.29\ Å$

$\pi \cdot r / 4 = 3.1\bar{4}\ (26.29)/4 = 20.65\ Å$

**1J8H-Vβ**

$$\iiint_V dV = \int_0^{70.38}\int_0^{20.65}\int_0^{27.91} \rho^2 \sin\phi\ d\rho\ d\theta\ d\phi \quad = \int_0^{70.38} 20.65\left[\frac{(26.29)^3}{3}\sin\phi\right] d\phi \quad \text{where, } \int \sin\phi = -\cos\phi$$

$$= \int_0^{70.38}\int_0^{20.65} \frac{1}{3}\rho^3 \sin\phi \Big|_{\rho=0}^{\rho=26.29} d\theta\ d\phi \quad = -20.65\left[\frac{(26.29)^3}{3}\right]\cos\phi \Big|_{\phi=0}^{\phi=70.38} \quad \text{where, } \cos 0 = 1$$

$$= \int_0^{70.38}\int_0^{20.65} \frac{(26.29)^3}{3}\sin\phi\ d\theta\ d\phi \quad = .336\left[-20.65\left[\frac{(26.29)^3}{3}\right]\right] - 1\left[-20.65\left[\frac{(26.29)^3}{3}\right]\right]$$

$$= \int_0^{70.38}\left[\frac{(26.29)^3}{3}\sin\phi\right]\theta \Big|_{\theta=0}^{\theta=20.65} d\phi \quad = -50{,}283 - \left[-149{,}651\right] = 99{,}368 \quad \text{where, multipl. x } \frac{\pi}{180} \text{ corrects for angular integrand of } \phi$$

$$\therefore \iiint_V dV = 1{,}734.30\ Å^3$$

**3T0E-Vα**

$\lambda = 132.31^\circ$ $\rho = 29.99\,\text{Å}$

$\phi = 180 - 132.31 = 47.69^\circ$ $r = \sin\phi\,(\rho) = 0.708\,(29.99) = 21.23\,\text{Å}$

$\pi \cdot r/4 = 3.1\bar{4}\,(21.23)/4 = 16.67\,\text{Å}$

$$\iiint_V dV = \int_0^{47.69}\int_0^{16.67}\int_0^{29.99} \rho^2 \sin\phi \; d\rho\, d\theta\, d\phi$$

$$= \int_0^{47.69}\int_0^{16.67} \frac{1}{3}\rho^3 \sin\phi \Big|_{\rho=0}^{\rho=29.99} d\theta\, d\phi$$

$$= \int_0^{47.69}\int_0^{16.67} \frac{(29.99)^3}{3}\sin\phi \; d\theta\, d\phi$$

$$= \int_0^{47.69} \left[\frac{(29.99)^3}{3}\sin\phi\right]\theta \Big|_{\theta=0}^{\theta=16.67} d\phi$$

$$= \int_0^{47.69} 16.67\left[\frac{(29.99)^3}{3}\sin\phi\right] d\phi \qquad \text{where, } \int \sin\phi = -\cos\phi$$

$$= -16.67\left[\frac{(29.99)^3}{3}\right]\cos\phi \Big|_{\phi=0}^{\phi=47.69} \qquad \text{where, } \cos 0 = 1$$

$$= .673\left[-16.67\left[\frac{(29.99)^3}{3}\right]\right] - 1\left[-16.67\left[\frac{(29.99)^3}{3}\right]\right]$$

$$= -100,869 - \left[-149,880\right] = 49,011 \qquad \text{where, multipl. x } \frac{\pi}{180}$$

corrects for angular integrand of $\phi$

$$\therefore \iiint_V dV = 855.40\ \text{Å}^3$$

**3T0E-Vβ**

$\lambda = 110.49^\circ$ $\rho = 24.63\,\text{Å}$

$\phi = 180 - 110.49 = 69.51^\circ$ $r = \sin\phi\,(\rho) = 0.937\,(24.63) = 23.08\,\text{Å}$

$\pi \cdot r/4 = 3.1\bar{4}\,(23.08)/4 = 18.13\,\text{Å}$

$$\iiint_V dV = \int_0^{69.51}\int_0^{18.13}\int_0^{24.63} \rho^2 \sin\phi \; d\rho\, d\theta\, d\phi$$

$$= \int_0^{69.51}\int_0^{18.13} \frac{1}{3}\rho^3 \sin\phi \Big|_{\rho=0}^{\rho=24.63} d\theta\, d\phi$$

$$= \int_0^{69.51}\int_0^{18.13} \frac{(24.63)^3}{3}\sin\phi \; d\theta\, d\phi$$

$$= \int_0^{69.51} \left[\frac{(24.63)^3}{3}\sin\phi\right]\theta \Big|_{\theta=0}^{\theta=18.13} d\phi$$

$$= \int_0^{69.51} 18.13\left[\frac{(24.63)^3}{3}\sin\phi\right] d\phi \qquad \text{where, } \int \sin\phi = -\cos\phi$$

$$= -18.13\left[\frac{(24.63)^3}{3}\right]\cos\phi \Big|_{\phi=0}^{\phi=69.51} \qquad \text{where, } \cos 0 = 1$$

$$= .350\left[-18.13\left[\frac{(24.63)^3}{3}\right]\right] - 1\left[-18.13\left[\frac{(24.63)^3}{3}\right]\right]$$

$$= -31,604 - \left[-90,296\right] = 58,692 \qquad \text{where, multipl. x } \frac{\pi}{180}$$

corrects for angular integrand of $\phi$

$$\therefore \iiint_V dV = 1,024.37\ \text{Å}^3$$

**1FYT-Vα**

$\lambda = 133.00^\circ$ $\rho = 29.70\,\text{Å}$

$\phi = 180 - 133.00 = 47.00^\circ$ $r = \sin\phi\,(\rho) = 0.731\,(29.70) = 21.71\,\text{Å}$

$\pi \cdot r/4 = 3.1\bar{4}\,(21.71)/4 = 17.05\,\text{Å}$

$$\iiint_V dV = \int_0^{47.00}\int_0^{17.05}\int_0^{29.70} \rho^2 \sin\phi \; d\rho\, d\theta\, d\phi$$

$$= \int_0^{47.00}\int_0^{17.05} \frac{1}{3}\rho^3 \sin\phi \Big|_{\rho=0}^{\rho=29.70} d\theta\, d\phi$$

$$= \int_0^{47.00}\int_0^{17.05} \frac{(29.70)^3}{3}\sin\phi \; d\theta\, d\phi$$

$$= \int_0^{47.00} \left[\frac{(29.70)^3}{3}\sin\phi\right]\theta \Big|_{\theta=0}^{\theta=17.05} d\phi$$

$$= \int_0^{47.00} 17.05\left[\frac{(29.70)^3}{3}\sin\phi\right] d\phi \qquad \text{where, } \int \sin\phi = -\cos\phi$$

$$= -17.05\left[\frac{(29.70)^3}{3}\right]\cos\phi \Big|_{\phi=0}^{\phi=47.00} \qquad \text{where, } \cos 0 = 1$$

$$= .682\left[-17.05\left[\frac{(29.70)^3}{3}\right]\right] - 1\left[-17.05\left[\frac{(29.70)^3}{3}\right]\right]$$

$$= -101,545 - \left[-148,892\right] = 47,347 \qquad \text{where, multipl. x } \frac{\pi}{180}$$

corrects for angular integrand of $\phi$

$$\therefore \iiint_V dV = 826.36\ \text{Å}^3$$

**1FYT-Vβ**

$\lambda = 109.97^\circ$ $\rho = 28.04\,\text{Å}$

$\phi = 180 - 109.97 = 70.03^\circ$ $r = \sin\phi\,(\rho) = 0.940\,(28.04) = 26.36\,\text{Å}$

$\pi \cdot r/4 = 3.1\bar{4}\,(26.36)/4 = 20.70\,\text{Å}$

$$\iiint_V dV = \int_0^{70.03}\int_0^{20.70}\int_0^{28.04} \rho^2 \sin\phi \; d\rho\, d\theta\, d\phi$$

$$= \int_0^{70.03}\int_0^{20.70} \frac{1}{3}\rho^3 \sin\phi \Big|_{\rho=0}^{\rho=28.04} d\theta\, d\phi$$

$$= \int_0^{70.03}\int_0^{20.70} \frac{(28.04)^3}{3}\sin\phi \; d\theta\, d\phi$$

$$= \int_0^{70.03} \left[\frac{(28.04)^3}{3}\sin\phi\right]\theta \Big|_{\theta=0}^{\theta=20.70} d\phi$$

$$= \int_0^{70.03} 20.70\left[\frac{(28.04)^3}{3}\sin\phi\right] d\phi \qquad \text{where, } \int \sin\phi = -\cos\phi$$

$$= -20.70\left[\frac{(28.04)^3}{3}\right]\cos\phi \Big|_{\phi=0}^{\phi=70.03} \qquad \text{where, } \cos 0 = 1$$

$$= .342\left[-20.70\left[\frac{(28.04)^3}{3}\right]\right] - 1\left[-20.70\left[\frac{(28.04)^3}{3}\right]\right]$$

$$= -52,025 - \left[-152,119\right] = 100,094 \qquad \text{where, multipl. x } \frac{\pi}{180}$$

corrects for angular integrand of $\phi$

$$\therefore \iiint_V dV = 1,746.97\ \text{Å}^3$$

**4H1L-Vα**

$\lambda = 125.30^\circ \qquad \rho = 32.62\ \text{Å}$

$\phi = 180 - 125.30 = 54.70^\circ \qquad r = \sin\phi\,(\rho) = 0.816(32.62) = 26.62\ \text{Å}$

$\pi \cdot r/4 = 3.1\bar{4}(26.62)/4 = 20.91\ \text{Å}$

$$\iiint_V dV = \int_0^{54.70}\int_0^{20.91}\int_0^{32.62} \rho^2 \sin\phi \; d\rho\, d\theta\, d\phi$$

$$= \int_0^{54.70}\int_0^{20.91} \frac{1}{3}\rho^3 \sin\phi \Big|_{\rho=0}^{\rho=32.62} d\theta\, d\phi$$

$$= \int_0^{54.70}\int_0^{20.91} \frac{(32.62)^3}{3}\sin\phi \; d\theta\, d\phi$$

$$= \int_0^{54.70} \left[\frac{(32.62)^3}{3}\sin\phi\right]\theta \Big|_{\theta=0}^{\theta=20.91} d\phi$$

$$= \int_0^{54.70} 20.91\left[\frac{(32.62)^3}{3}\sin\phi\right] d\phi \qquad \text{where, } \int \sin\phi = -\cos\phi$$

$$= -\,20.91\left[\frac{(32.62)^3}{3}\right]\cos\phi \Big|_{\phi=0}^{\phi=54.70} \qquad \text{where, } \cos 0 = 1$$

$$= .578\left[-\,20.91\left[\frac{(32.62)^3}{3}\right]\right] - 1\left[-20.91\left[\frac{(32.62)^3}{3}\right]\right]$$

$$= -139,834 - \left[-\,241,927\right] = 102,093 \qquad \text{where, multipl. x } \frac{\pi}{180} \text{ corrects for angular integrand of } \phi$$

$$\therefore \iiint_V dV = 1,781.86\ \text{Å}^3$$

**4H1L-Vβ**

$\lambda = 117.27^\circ \qquad \rho = 27.16\ \text{Å}$

$\phi = 180 - 117.27 = 62.73^\circ \qquad r = \sin\phi\,(\rho) = 0.889(27.16) = 24.15\ \text{Å}$

$\pi \cdot r/4 = 3.1\bar{4}(24.15)/4 = 18.97\ \text{Å}$

$$\iiint_V dV = \int_0^{62.73}\int_0^{18.97}\int_0^{27.16} \rho^2 \sin\phi \; d\rho\, d\theta\, d\phi$$

$$= \int_0^{62.73}\int_0^{18.97} \frac{1}{3}\rho^3 \sin\phi \Big|_{\rho=0}^{\rho=27.16} d\theta\, d\phi$$

$$= \int_0^{62.73}\int_0^{18.97} \frac{(27.16)^3}{3}\sin\phi \; d\theta\, d\phi$$

$$= \int_0^{62.73} \left[\frac{(27.16)^3}{3}\sin\phi\right]\theta \Big|_{\theta=0}^{\theta=18.97} d\phi$$

$$= \int_0^{62.73} 18.97\left[\frac{(27.16)^3}{3}\sin\phi\right] d\phi \qquad \text{where, } \int \sin\phi = -\cos\phi$$

$$= -\,18.97\left[\frac{(27.16)^3}{3}\right]\cos\phi \Big|_{\phi=0}^{\phi=62.73} \qquad \text{where, } \cos 0 = 1$$

$$= .458\left[-\,18.97\left[\frac{(27.16)^3}{3}\right]\right] - 1\left[-18.97\left[\frac{(27.16)^3}{3}\right]\right]$$

$$= -\,58,023 - \left[-126,688\right] = 68,665 \qquad \text{where, multipl. x } \frac{\pi}{180} \text{ corrects for angular integrand of } \phi$$

$$\therefore \iiint_V dV = 1,198.43\ \text{Å}^3$$

**1ZGL-Vα**

$\lambda = 117.60^\circ \qquad \rho = 33.78\ \text{Å}$

$\phi = 180 - 117.60 = 62.40^\circ \qquad r = \sin\phi\,(\rho) = 0.886(33.78) = 29.93\ \text{Å}$

$\pi \cdot r/4 = 3.1\bar{4}(29.93)/4 = 23.51\ \text{Å}$

$$\iiint_V dV = \int_0^{62.40}\int_0^{23.51}\int_0^{33.78} \rho^2 \sin\phi \; d\rho\, d\theta\, d\phi$$

$$= \int_0^{62.40}\int_0^{23.51} \frac{1}{3}\rho^3 \sin\phi \Big|_{\rho=0}^{\rho=33.78} d\theta\, d\phi$$

$$= \int_0^{62.40}\int_0^{23.51} \frac{(33.78)^3}{3}\sin\phi \; d\theta\, d\phi$$

$$= \int_0^{62.40} \left[\frac{(33.78)^3}{3}\sin\phi\right]\theta \Big|_{\theta=0}^{\theta=23.51} d\phi$$

$$= \int_0^{62.40} 23.51\left[\frac{(33.78)^3}{3}\sin\phi\right] d\phi \qquad \text{where, } \int \sin\phi = -\cos\phi$$

$$= -\,23.51\left[\frac{(33.78)^3}{3}\right]\cos\phi \Big|_{\phi=0}^{\phi=62.40} \qquad \text{where, } \cos 0 = 1$$

$$= .463\left[-\,23.51\left[\frac{(33.78)^3}{3}\right]\right] - 1\left[-23.51\left[\frac{(33.78)^3}{3}\right]\right]$$

$$= -139,859 - \left[-\,302,072\right] = 162,213 \qquad \text{where, multipl. x } \frac{\pi}{180} \text{ corrects for angular integrand of } \phi$$

$$\therefore \iiint_V dV = 2,831.15\ \text{Å}^3$$

**1ZGL-Vβ**

$\lambda = 108.58^\circ \qquad \rho = 25.67\ \text{Å}$

$\phi = 180 - 108.58 = 71.42^\circ \qquad r = \sin\phi\,(\rho) = 0.948(25.67) = 24.33\ \text{Å}$

$\pi \cdot r/4 = 3.1\bar{4}(26.36)/4 = 19.11\ \text{Å}$

$$\iiint_V dV = \int_0^{71.42}\int_0^{19.11}\int_0^{25.67} \rho^2 \sin\phi \; d\rho\, d\theta\, d\phi$$

$$= \int_0^{71.42}\int_0^{19.11} \frac{1}{3}\rho^3 \sin\phi \Big|_{\rho=0}^{\rho=25.67} d\theta\, d\phi$$

$$= \int_0^{71.42}\int_0^{19.11} \frac{(25.67)^3}{3}\sin\phi \; d\theta\, d\phi$$

$$= \int_0^{71.42} \left[\frac{(25.67)^3}{3}\sin\phi\right]\theta \Big|_{\theta=0}^{\theta=19.11} d\phi$$

$$= \int_0^{71.42} 19.11\left[\frac{(25.67)^3}{3}\sin\phi\right] d\phi \qquad \text{where, } \int \sin\phi = -\cos\phi$$

$$= -19.11\left[\frac{(25.67)^3}{3}\right]\cos\phi \Big|_{\phi=0}^{\phi=71.42} \qquad \text{where, } \cos 0 = 1$$

$$= .319\left[-19.11\left[\frac{(25.67)^3}{3}\right]\right] - 1\left[-19.11\left[\frac{(25.67)^3}{3}\right]\right]$$

$$= -\,34,372 - \left[-107,750\right] = 73,378 \qquad \text{where, multipl. x } \frac{\pi}{180} \text{ corrects for angular integrand of } \phi$$

$$\therefore \iiint_V dV = 1,280.68\ \text{Å}^3$$

##### 2IAN-Vα

$\lambda = 116.16°$ $\rho = 29.39\,\text{Å}$

$\phi = 180 - 116.16 = 63.84°$ $r = \sin\phi\,(\rho) = 0.898\,(29.39) = 26.38\,\text{Å}$

$\pi \cdot r/4 = 3.1\bar{4}\,(26.38)/4 = 20.71\,\text{Å}$

$$\iiint_V dV = \int_0^{63.84}\int_0^{20.71}\int_0^{29.39} \rho^2 \sin\phi \; d\rho\; d\theta\; d\phi$$

$$= \int_0^{63.84}\int_0^{20.71} \frac{1}{3}\rho^3 \sin\phi \Big|_{\rho=0}^{\rho=29.39} d\theta\; d\phi$$

$$= \int_0^{63.84}\int_0^{20.71} \frac{(29.39)^3}{3}\sin\phi \; d\theta\; d\phi$$

$$= \int_0^{63.84} \left[\frac{(29.39)^3}{3}\sin\phi\right]\theta \Big|_{\theta=0}^{\theta=20.71} d\phi$$

$$= \int_0^{63.84} 20.71\left[\frac{(29.39)^3}{3}\sin\phi\right] d\phi \qquad \text{where, } \int \sin\phi = -\cos\phi$$

$$= -20.71\left[\frac{(29.39)^3}{3}\right]\cos\phi \Big|_{\phi=0}^{\phi=63.84} \qquad \text{where, } \cos 0 = 1$$

$$= .441\left[-20.71\left[\frac{(29.39)^3}{3}\right]\right] - 1\left[-20.71\left[\frac{(29.39)^3}{3}\right]\right]$$

$$= -77,285 - \left[-175,250\right] = 97,965 \qquad \text{where, multipl. x } \frac{\pi}{180} \text{ corrects for angular integrand of } \phi$$

$$\therefore \iiint_V dV = 1,709.81\,\text{Å}^3$$

##### 2IAN-Vβ

$\lambda = 130.76°$ $\rho = 21.78\,\text{Å}$

$\phi = 180 - 130.76 = 49.24°$ $r = \sin\phi\,(\rho) = 0.757\,(21.78) = 16.50\,\text{Å}$

$\pi \cdot r/4 = 3.1\bar{4}\,(16.50)/4 = 12.96\,\text{Å}$

$$\iiint_V dV = \int_0^{49.24}\int_0^{12.96}\int_0^{21.78} \rho^2 \sin\phi \; d\rho\; d\theta\; d\phi$$

$$= \int_0^{49.24}\int_0^{12.96} \frac{1}{3}\rho^3 \sin\phi \Big|_{\rho=0}^{\rho=21.78} d\theta\; d\phi$$

$$= \int_0^{49.24}\int_0^{12.96} \frac{(21.78)^3}{3}\sin\phi \; d\theta\; d\phi$$

$$= \int_0^{49.24} \left[\frac{(21.78)^3}{3}\sin\phi\right]\theta \Big|_{\theta=0}^{\theta=12.96} d\phi$$

$$= \int_0^{49.24} 12.96\left[\frac{(21.78)^3}{3}\sin\phi\right] d\phi \qquad \text{where, } \int \sin\phi = -\cos\phi$$

$$= -12.96\left[\frac{(21.78)^3}{3}\right]\cos\phi \Big|_{\phi=0}^{\phi=49.24} \qquad \text{where, } \cos 0 = 1$$

$$= .653\left[-12.96\left[\frac{(21.78)^3}{3}\right]\right] - 1\left[-12.96\left[\frac{(21.78)^3}{3}\right]\right]$$

$$= -29,145 - \left[-44,633\right] = 15,488 \qquad \text{where, multipl. x } \frac{\pi}{180} \text{ corrects for angular integrand of } \phi$$

$$\therefore \iiint_V dV = 270.31\,\text{Å}^3$$

### 4E41-Vα

$\lambda = 119.79°$ $\rho = 29.51\,\text{Å}$

$\phi = 180 - 119.79 = 60.21°$ $r = \sin\phi\,(\rho) = 0.868\,(29.51) = 25.61\,\text{Å}$

$\pi \cdot r/4 = 3.1\bar{4}\,(25.61)/4 = 20.11\,\text{Å}$

$$\iiint_V dV = \int_0^{60.21}\int_0^{20.11}\int_0^{29.51} \rho^2 \sin\phi \; d\rho\; d\theta\; d\phi$$

$$= \int_0^{60.21}\int_0^{20.11} \frac{1}{3}\rho^3 \sin\phi \Big|_{\rho=0}^{\rho=29.51} d\theta\; d\phi$$

$$= \int_0^{60.21}\int_0^{20.11} \frac{(29.51)^3}{3}\sin\phi \; d\theta\; d\phi$$

$$= \int_0^{60.21} \left[\frac{(29.51)^3}{3}\sin\phi\right]\theta \Big|_{\theta=0}^{\theta=20.11} d\phi$$

$$= \int_0^{60.21} 20.11\left[\frac{(29.51)^3}{3}\sin\phi\right] d\phi \qquad \text{where, } \int \sin\phi = -\cos\phi$$

$$= -20.11\left[\frac{(29.51)^3}{3}\right]\cos\phi \Big|_{\phi=0}^{\phi=60.21} \qquad \text{where, } \cos 0 = 1$$

$$= .497\left[-20.11\left[\frac{(29.51)^3}{3}\right]\right] - 1\left[-20.11\left[\frac{(29.51)^3}{3}\right]\right]$$

$$= -85,616 - \left[-172,266\right] = 86,650 \qquad \text{where, multipl. x } \frac{\pi}{180} \text{ corrects for angular integrand of } \phi$$

$$\therefore \iiint_V dV = 1,512.33\,\text{Å}^3$$

### 4E41-Vβ

$\lambda = 129.37°$ $\rho = 23.67\,\text{Å}$

$\phi = 180 - 129.37 = 50.63°$ $r = \sin\phi\,(\rho) = 0.773\,(23.67) = 18.30\,\text{Å}$

$\pi \cdot r/4 = 3.1\bar{4}\,(18.30)/4 = 14.37\,\text{Å}$

$$\iiint_V dV = \int_0^{50.63}\int_0^{14.37}\int_0^{23.67} \rho^2 \sin\phi \; d\rho\; d\theta\; d\phi$$

$$= \int_0^{50.63}\int_0^{14.37} \frac{1}{3}\rho^3 \sin\phi \Big|_{\rho=0}^{\rho=23.67} d\theta\; d\phi$$

$$= \int_0^{50.63}\int_0^{14.37} \frac{(23.67)^3}{3}\sin\phi \; d\theta\; d\phi$$

$$= \int_0^{50.63} \left[\frac{(23.67)^3}{3}\sin\phi\right]\theta \Big|_{\theta=0}^{\theta=14.37} d\phi$$

$$= \int_0^{50.63} 14.37\left[\frac{(23.67)^3}{3}\sin\phi\right] d\phi \qquad \text{where, } \int \sin\phi = -\cos\phi$$

$$= -14.37\left[\frac{(23.67)^3}{3}\right]\cos\phi \Big|_{\phi=0}^{\phi=50.63} \qquad \text{where, } \cos 0 = 1$$

$$= .634\left[-14.37\left[\frac{(23.67)^3}{3}\right]\right] - 1\left[-14.37\left[\frac{(23.67)^3}{3}\right]\right]$$

$$= -40,274 - \left[-63,523\right] = 23,249 \qquad \text{where, multipl. x } \frac{\pi}{180} \text{ corrects for angular integrand of } \phi$$

$$\therefore \iiint_V dV = 405.77\,\text{Å}^3$$

6CQL-Vα

$\lambda = 133.17^\circ$ $\rho = 29.01\,\mathring{A}$

$\phi = 180 - 133.17 = 46.83^\circ$ $r = \sin\phi\,(\rho) = 0.729\,(29.01) = 21.16\,\mathring{A}$

$\pi \cdot r/4 = 3.1\bar{4}\,(21.16)/4 = 16.62\,\mathring{A}$

$$\iiint_V dV = \int_0^{46.83}\int_0^{16.62}\int_0^{29.01} \rho^2 \sin\phi \; d\rho\, d\theta\, d\phi$$

$$= \int_0^{46.83}\int_0^{16.62} \frac{1}{3}\rho^3 \sin\phi \Big|_{\rho=0}^{\rho=29.01} d\theta\, d\phi$$

$$= \int_0^{46.83}\int_0^{16.62} \frac{(29.01)^3}{3} \sin\phi \; d\theta\, d\phi$$

$$= \int_0^{46.83} \left[\frac{(29.01)^3}{3}\sin\phi\right] \theta \Big|_{\theta=0}^{\theta=16.62} d\phi$$

$$= \int_0^{46.83} 16.62 \left[\frac{(29.01)^3}{3}\sin\phi\right] d\phi \qquad \text{where, } \int \sin\phi = -\cos\phi$$

$$= -16.62\left[\frac{(29.01)^3}{3}\right]\cos\phi \Big|_{\phi=0}^{\phi=46.83} \qquad \text{where, } \cos 0 = 1$$

$$= .684\left[-16.62\left[\frac{(29.01)^3}{3}\right]\right] - 1\left[-16.62\left[\frac{(29.01)^3}{3}\right]\right]$$

$$= -92,514 - \left[-135,255\right] = 42,741 \qquad \text{where, multipl. x } \frac{\pi}{180} \text{ corrects for angular integrand of } \phi$$

$$\therefore \iiint_V dV = 745.97\,\mathring{A}^3$$

6CQL-Vβ

$\lambda = 129.70^\circ$ $\rho = 25.56\,\mathring{A}$

$\phi = 180 - 129.70 = 50.30^\circ$ $r = \sin\phi\,(\rho) = 0.769\,(25.56) = 19.67\,\mathring{A}$

$\pi \cdot r/4 = 3.1\bar{4}\,(19.67)/4 = 15.45\,\mathring{A}$

$$\iiint_V dV = \int_0^{50.30}\int_0^{15.45}\int_0^{25.56} \rho^2 \sin\phi \; d\rho\, d\theta\, d\phi$$

$$= \int_0^{50.30}\int_0^{15.45} \frac{1}{3}\rho^3 \sin\phi \Big|_{\rho=0}^{\rho=25.56} d\theta\, d\phi$$

$$= \int_0^{50.30}\int_0^{15.45} \frac{(25.56)^3}{3} \sin\phi \; d\theta\, d\phi$$

$$= \int_0^{50.30} \left[\frac{(25.56)^3}{3}\sin\phi\right] \theta \Big|_{\theta=0}^{\theta=15.45} d\phi$$

$$= \int_0^{50.30} 15.45 \left[\frac{(25.56)^3}{3}\sin\phi\right] d\phi \qquad \text{where, } \int \sin\phi = -\cos\phi$$

$$= -15.45\left[\frac{(25.56)^3}{3}\right]\cos\phi \Big|_{\phi=0}^{\phi=50.30} \qquad \text{where, } \cos 0 = 1$$

$$= .639\left[-15.45\left[\frac{(25.56)^3}{3}\right]\right] - 1\left[-15.45\left[\frac{(25.56)^3}{3}\right]\right]$$

$$= -54,953 - \left[-85,998\right] = 31,045 \qquad \text{where, multipl. x } \frac{\pi}{180} \text{ corrects for angular integrand of } \phi$$

$$\therefore \iiint_V dV = 541.84\,\mathring{A}^3$$

6CQN-Vα

$\lambda = 134.45^\circ$ $\rho = 28.98\,\mathring{A}$

$\phi = 180 - 134.45 = 45.55^\circ$ $r = \sin\phi\,(\rho) = 0.714\,(28.98) = 20.69\,\mathring{A}$

$\pi \cdot r/4 = 3.1\bar{4}\,(20.69)/4 = 16.25\,\mathring{A}$

$$\iiint_V dV = \int_0^{45.55}\int_0^{16.25}\int_0^{28.98} \rho^2 \sin\phi \; d\rho\, d\theta\, d\phi$$

$$= \int_0^{45.55}\int_0^{16.25} \frac{1}{3}\rho^3 \sin\phi \Big|_{\rho=0}^{\rho=28.98} d\theta\, d\phi$$

$$= \int_0^{45.55}\int_0^{16.25} \frac{(28.98)^3}{3} \sin\phi \; d\theta\, d\phi$$

$$= \int_0^{45.55} \left[\frac{(28.98)^3}{3}\sin\phi\right] \theta \Big|_{\theta=0}^{\theta=16.25} d\phi$$

$$= \int_0^{45.55} 16.25 \left[\frac{(28.98)^3}{3}\sin\phi\right] d\phi \qquad \text{where, } \int \sin\phi = -\cos\phi$$

$$= -16.25\left[\frac{(28.98)^3}{3}\right]\cos\phi \Big|_{\phi=0}^{\phi=45.55} \qquad \text{where, } \cos 0 = 1$$

$$= .700\left[-16.25\left[\frac{(28.98)^3}{3}\right]\right] - 1\left[-16.25\left[\frac{(28.98)^3}{3}\right]\right]$$

$$= -92,284 - \left[-131,834\right] = 39,550 \qquad \text{where, multipl. x } \frac{\pi}{180} \text{ corrects for angular integrand of } \phi$$

$$\therefore \iiint_V dV = 690.28\,\mathring{A}^3$$

6CQN-Vβ

$\lambda = 128.57^\circ$ $\rho = 25.68\,\mathring{A}$

$\phi = 180 - 128.57 = 51.43^\circ$ $r = \sin\phi\,(\rho) = 0.782\,(25.68) = 20.08\,\mathring{A}$

$\pi \cdot r/4 = 3.1\bar{4}\,(20.08)/4 = 15.77\,\mathring{A}$

$$\iiint_V dV = \int_0^{51.43}\int_0^{15.77}\int_0^{25.68} \rho^2 \sin\phi \; d\rho\, d\theta\, d\phi$$

$$= \int_0^{51.43}\int_0^{15.77} \frac{1}{3}\rho^3 \sin\phi \Big|_{\rho=0}^{\rho=25.68} d\theta\, d\phi$$

$$= \int_0^{51.43}\int_0^{15.77} \frac{(25.68)^3}{3} \sin\phi \; d\theta\, d\phi$$

$$= \int_0^{51.43} \left[\frac{(25.68)^3}{3}\sin\phi\right] \theta \Big|_{\theta=0}^{\theta=15.77} d\phi$$

$$= \int_0^{51.43} 15.77 \left[\frac{(25.68)^3}{3}\sin\phi\right] d\phi \qquad \text{where, } \int \sin\phi = -\cos\phi$$

$$= -15.77\left[\frac{(25.68)^3}{3}\right]\cos\phi \Big|_{\phi=0}^{\phi=51.43} \qquad \text{where, } \cos 0 = 1$$

$$= .623\left[-15.77\left[\frac{(25.68)^3}{3}\right]\right] - 1\left[-15.77\left[\frac{(25.68)^3}{3}\right]\right]$$

$$= -55,460 - \left[-89,022\right] = 33,562 \qquad \text{where, multipl. x } \frac{\pi}{180} \text{ corrects for angular integrand of } \phi$$

$$\therefore \iiint_V dV = 585.76\,\mathring{A}^3$$

$\lambda = 134.32^\circ$ $\rho = 28.97\,\mathring{A}$

$\phi = 180 - 134.32 = 45.68^\circ$ $r = \sin\phi\,(\rho) = 0.715\,(28.97) = 20.73\,\mathring{A}$

$\pi \cdot r/4 = 3.14\,(20.73)/4 = 16.28\,\mathring{A}$

6CQQ-Vα

$$\iiint_V dV = \int_0^{45.68}\int_0^{16.28}\int_0^{28.97} \rho^2 \sin\phi \; d\rho \; d\theta \; d\phi$$

$$= \int_0^{45.68}\int_0^{16.28} \tfrac{1}{3}\rho^3 \sin\phi \Big|_{\rho=0}^{\rho=28.97} d\theta \; d\phi$$

$$= \int_0^{45.68}\int_0^{16.28} \frac{(28.97)^3}{3}\sin\phi \; d\theta \; d\phi$$

$$= \int_0^{45.68} \left[\frac{(28.97)^3}{3}\sin\phi\right]\theta \Big|_{\theta=0}^{\theta=16.28} d\phi$$

$$= \int_0^{45.68} 16.28\left[\frac{(28.97)^3}{3}\sin\phi\right] d\phi \qquad \text{where, } \int \sin\phi = -\cos\phi$$

$$= -16.28\left[\frac{(28.97)^3}{3}\right]\cos\phi \Big|_{\phi=0}^{\phi=45.68} \qquad \text{where, } \cos 0 = 1$$

$$= .699\left[-16.28\left[\frac{(28.97)^3}{3}\right]\right] - 1\left[-16.28\left[\frac{(28.97)^3}{3}\right]\right]$$

$$= -92,227 - \left[-131,941\right] = 39,714 \qquad \text{where, multipl. x } \frac{\pi}{180} \text{ corrects for angular integrand of } \phi$$

$$\therefore \iiint_V dV = 693.14\,\mathring{A}^3$$

$\lambda = 127.12^\circ$ $\rho = 26.34\,\mathring{A}$

$\phi = 180 - 127.12 = 55.90^\circ$ $r = \sin\phi\,(\rho) = 0.828\,(26.34) = 21.81\,\mathring{A}$

$\pi \cdot r/4 = 3.14\,(21.81)/4 = 17.13\,\mathring{A}$

6CQQ-Vβ

$$\iiint_V dV = \int_0^{55.90}\int_0^{17.13}\int_0^{26.34} \rho^2 \sin\phi \; d\rho \; d\theta \; d\phi$$

$$= \int_0^{55.90}\int_0^{17.13} \tfrac{1}{3}\rho^3 \sin\phi \Big|_{\rho=0}^{\rho=26.34} d\theta \; d\phi$$

$$= \int_0^{55.90}\int_0^{17.13} \frac{(26.34)^3}{3}\sin\phi \; d\theta \; d\phi$$

$$= \int_0^{55.90} \left[\frac{(26.34)^3}{3}\sin\phi\right]\theta \Big|_{\theta=0}^{\theta=17.13} d\phi$$

$$= \int_0^{55.90} 17.13\left[\frac{(26.34)^3}{3}\sin\phi\right] d\phi \qquad \text{where, } \int \sin\phi = -\cos\phi$$

$$= -17.13\left[\frac{(26.34)^3}{3}\right]\cos\phi \Big|_{\phi=0}^{\phi=55.90} \qquad \text{where, } \cos 0 = 1$$

$$= .561\left[-17.13\left[\frac{(26.34)^3}{3}\right]\right] - 1\left[-17.13\left[\frac{(26.34)^3}{3}\right]\right]$$

$$= -58,539 - \left[-104,348\right] = 45,809 \qquad \text{where, multipl. x } \frac{\pi}{180} \text{ corrects for angular integrand of } \phi$$

$$\therefore \iiint_V dV = 799.51\,\mathring{A}^3$$

$\lambda = 133.75^\circ$ $\rho = 28.99\,\mathring{A}$

$\phi = 180 - 133.75 = 46.25^\circ$ $r = \sin\phi\,(\rho) = 0.722(28.99) = 20.94\,\mathring{A}$

$\pi \cdot r/4 = 3.14\,(20.94)/4 = 16.45\,\mathring{A}$

6CQR-Vα

$$\iiint_V dV = \int_0^{46.25}\int_0^{16.45}\int_0^{28.99} \rho^2 \sin\phi \; d\rho \; d\theta \; d\phi$$

$$= \int_0^{46.25}\int_0^{16.45} \tfrac{1}{3}\rho^3 \sin\phi \Big|_{\rho=0}^{\rho=28.99} d\theta \; d\phi$$

$$= \int_0^{46.25}\int_0^{16.45} \frac{(28.99)^3}{3}\sin\phi \; d\theta \; d\phi$$

$$= \int_0^{46.25} \left[\frac{(28.99)^3}{3}\sin\phi\right]\theta \Big|_{\theta=0}^{\theta=16.45} d\phi$$

$$= \int_0^{46.25} 16.45\left[\frac{(28.99)^3}{3}\sin\phi\right] d\phi \qquad \text{where, } \int \sin\phi = -\cos\phi$$

$$= -16.45\left[\frac{(28.99)^3}{3}\right]\cos\phi \Big|_{\phi=0}^{\phi=46.25} \qquad \text{where, } \cos 0 = 1$$

$$= .692\left[-16.45\left[\frac{(28.99)^3}{3}\right]\right] - 1\left[-16.45\left[\frac{(28.99)^3}{3}\right]\right]$$

$$= -92,448 - \left[-133,595\right] = 41,147 \qquad \text{where, multipl. x } \frac{\pi}{180} \text{ corrects for angular integrand of } \phi$$

$$\therefore \iiint_V dV = 718.16\,\mathring{A}^3$$

$\lambda = 124.36^\circ$ $\rho = 25.95\,\mathring{A}$

$\phi = 180 - 124.36 = 55.64^\circ$ $r = \sin\phi\,(\rho) = 0.826(25.95) = 21.42\,\mathring{A}$

$\pi \cdot r/4 = 3.14\,(21.42)/4 = 16.82\,\mathring{A}$

6CQR-Vβ

$$\iiint_V dV = \int_0^{55.64}\int_0^{16.82}\int_0^{25.95} \rho^2 \sin\phi \; d\rho \; d\theta \; d\phi$$

$$= \int_0^{55.64}\int_0^{16.82} \tfrac{1}{3}\rho^3 \sin\phi \Big|_{\rho=0}^{\rho=25.95} d\theta \; d\phi$$

$$= \int_0^{55.64}\int_0^{16.82} \frac{(25.95)^3}{3}\sin\phi \; d\theta \; d\phi$$

$$= \int_0^{55.64} \left[\frac{(25.95)^3}{3}\sin\phi\right]\theta \Big|_{\theta=0}^{\theta=16.82} d\phi$$

$$= \int_0^{55.64} 16.82\left[\frac{(25.95)^3}{3}\sin\phi\right] d\phi \qquad \text{where, } \int \sin\phi = -\cos\phi$$

$$= -16.82\left[\frac{(25.95)^3}{3}\right]\cos\phi \Big|_{\phi=0}^{\phi=55.64} \qquad \text{where, } \cos 0 = 1$$

$$= .564\left[-16.82\left[\frac{(25.95)^3}{3}\right]\right] - 1\left[-16.82\left[\frac{(25.95)^3}{3}\right]\right]$$

$$= -55,258 - \left[-97,975\right] = 42,717 \qquad \text{where, multipl. x } \frac{\pi}{180} \text{ corrects for angular integrand of } \phi$$

$$\therefore \iiint_V dV = 745.55\,\mathring{A}^3$$

5KSA-Vα

$\lambda = 134.95^\circ$ $\rho = 28.76\ \text{Å}$

$\phi = 180 - 134.95 = 45.05^\circ$ $r = \sin\phi\,(\rho) = 0.708(28.76) = 20.35\ \text{Å}$

$\pi \cdot r / 4 = 3.1\bar{4}\,(20.35)/4 = 15.99\ \text{Å}$

$$\iiint_V dV = \int_0^{45.05}\int_0^{15.99}\int_0^{28.76} \rho^2 \sin\phi \, d\rho \, d\theta \, d\phi$$

$$= \int_0^{45.05}\int_0^{15.99} \frac{1}{3}\rho^3 \sin\phi \Big|_{\rho=0}^{\rho=28.76} d\theta \, d\phi$$

$$= \int_0^{45.05}\int_0^{15.99} \frac{(28.76)^3}{3} \sin\phi \, d\theta \, d\phi$$

$$= \int_0^{45.05} \left[\frac{(28.76)^3}{3}\sin\phi\right] \theta \Big|_{\theta=0}^{\theta=15.99} d\phi$$

$$= \int_0^{45.05} 15.99 \left[\frac{(28.76)^3}{3}\sin\phi\right] d\phi \qquad \text{where, } \int \sin\phi = -\cos\phi$$

$$= -15.99\left[\frac{(28.76)^3}{3}\right]\cos\phi \Big|_{\phi=0}^{\phi=45.05} \qquad \text{where, } \cos 0 = 1$$

$$= .706\left[-15.99\left[\frac{(28.76)^3}{3}\right]\right] - 1\left[-15.99\left[\frac{(28.76)^3}{3}\right]\right]$$

$$= -89{,}516 - \left[-126{,}793\right] = 37{,}277 \qquad \text{where, multipl. x } \frac{\pi}{180}$$

corrects for angular integrand of $\phi$

$$\therefore \iiint_V dV = 650.61\ \text{Å}^3$$

5KSA-Vβ

$\lambda = 119.95^\circ$ $\rho = 26.02\ \text{Å}$

$\phi = 180 - 119.95 = 60.85^\circ$ $r = \sin\phi\,(\rho) = 0.873(26.02) = 22.72\ \text{Å}$

$\pi \cdot r / 4 = 3.1\bar{4}\,(22.72)/4 = 17.85\ \text{Å}$

$$\iiint_V dV = \int_0^{60.85}\int_0^{17.85}\int_0^{26.02} \rho^2 \sin\phi \, d\rho \, d\theta \, d\phi$$

$$= \int_0^{60.85}\int_0^{17.85} \frac{1}{3}\rho^3 \sin\phi \Big|_{\rho=0}^{\rho=26.02} d\theta \, d\phi$$

$$= \int_0^{60.85}\int_0^{17.85} \frac{(26.02)^3}{3} \sin\phi \, d\theta \, d\phi$$

$$= \int_0^{60.85} \left[\frac{(26.02)^3}{3}\sin\phi\right] \theta \Big|_{\theta=0}^{\theta=17.85} d\phi$$

$$= \int_0^{60.85} 17.85 \left[\frac{(26.02)^3}{3}\sin\phi\right] d\phi \qquad \text{where, } \int \sin\phi = -\cos\phi$$

$$= -17.85\left[\frac{(26.02)^3}{3}\right]\cos\phi \Big|_{\phi=0}^{\phi=60.85} \qquad \text{where, } \cos 0 = 1$$

$$= .487\left[-17.85\left[\frac{(26.02)^3}{3}\right]\right] - 1\left[-17.85\left[\frac{(26.02)^3}{3}\right]\right]$$

$$= -51{,}047 - \left[-104{,}819\right] = 53{,}772 \qquad \text{where, multipl. x } \frac{\pi}{180}$$

corrects for angular integrand of $\phi$

$$\therefore \iiint_V dV = 938.50\ \text{Å}^3$$

5KS9-Vα

$\lambda = 128.32^\circ$ $\rho = 28.65\ \text{Å}$

$\phi = 180 - 128.32 = 51.68^\circ$ $r = \sin\phi\,(\rho) = 0.785(28.65) = 22.48\ \text{Å}$

$\pi \cdot r / 4 = 3.1\bar{4}\,(22.48)/4 = 17.65\ \text{Å}$

$$\iiint_V dV = \int_0^{51.68}\int_0^{17.65}\int_0^{28.65} \rho^2 \sin\phi \, d\rho \, d\theta \, d\phi$$

$$= \int_0^{51.68}\int_0^{17.65} \frac{1}{3}\rho^3 \sin\phi \Big|_{\rho=0}^{\rho=28.65} d\theta \, d\phi$$

$$= \int_0^{51.68}\int_0^{17.65} \frac{(28.65)^3}{3} \sin\phi \, d\theta \, d\phi$$

$$= \int_0^{51.68} \left[\frac{(28.65)^3}{3}\sin\phi\right] \theta \Big|_{\theta=0}^{\theta=17.65} d\phi$$

$$= \int_0^{51.68} 17.65 \left[\frac{(28.65)^3}{3}\sin\phi\right] d\phi \qquad \text{where, } \int \sin\phi = -\cos\phi$$

$$= -17.65\left[\frac{(28.65)^3}{3}\right]\cos\phi \Big|_{\phi=0}^{\phi=51.68} \qquad \text{where, } \cos 0 = 1$$

$$= .620\left[-17.65\left[\frac{(28.65)^3}{3}\right]\right] - 1\left[-17.65\left[\frac{(28.65)^3}{3}\right]\right]$$

$$= -85{,}788 - \left[-138{,}356\right] = 52{,}568 \qquad \text{where, multipl. x } \frac{\pi}{180}$$

corrects for angular integrand of $\phi$

$$\therefore \iiint_V dV = 917.61\ \text{Å}^3$$

5KS9-Vβ

$\lambda = 126.25^\circ$ $\rho = 23.57\ \text{Å}$

$\phi = 180 - 126.25 = 60.85^\circ$ $r = \sin\phi\,(\rho) = 0.852(32.35) = 22.72\ \text{Å}$

$\pi \cdot r / 4 = 3.1\bar{4}\,(22.72)/4 = 17.85\ \text{Å}$

$$\iiint_V dV = \int_0^{60.85}\int_0^{17.85}\int_0^{23.57} \rho^2 \sin\phi \, d\rho \, d\theta \, d\phi$$

$$= \int_0^{60.85}\int_0^{17.85} \frac{1}{3}\rho^3 \sin\phi \Big|_{\rho=0}^{\rho=23.57} d\theta \, d\phi$$

$$= \int_0^{60.85}\int_0^{17.85} \frac{(23.57)^3}{3} \sin\phi \, d\theta \, d\phi$$

$$= \int_0^{60.85} \left[\frac{(23.57)^3}{3}\sin\phi\right] \theta \Big|_{\theta=0}^{\theta=17.85} d\phi$$

$$= \int_0^{60.85} 17.85 \left[\frac{(23.57)^3}{3}\sin\phi\right] d\phi \qquad \text{where, } \int \sin\phi = -\cos\phi$$

$$= -17.85\left[\frac{(23.57)^3}{3}\right]\cos\phi \Big|_{\phi=0}^{\phi=60.85} \qquad \text{where, } \cos 0 = 1$$

$$= .487\left[-17.85\left[\frac{(23.57)^3}{3}\right]\right] - 1\left[-17.85\left[\frac{(23.57)^3}{3}\right]\right]$$

$$= -37{,}942 - \left[-77{,}910\right] = 39{,}968 \qquad \text{where, multipl. x } \frac{\pi}{180}$$

corrects for angular integrand of $\phi$

$$\therefore \iiint_V dV = 697.57\ \text{Å}^3$$

**4OZF-Vα**

$\lambda = 131.50°$ $\quad \rho = 33.17\,Å$

$\phi = 180 - 131.50 = 48.50°$ $\quad r = \sin\phi\ (\rho) = 0.749(33.17) = 24.84\,Å$

$\pi \cdot r/4 = 3.14\,(24.84)/4 = 19.51\,Å$

$$\iiint_V dV = \int_0^{48.50}\int_0^{19.51}\int_0^{33.17} \rho^2 \sin\phi\ d\rho\ d\theta\ d\phi$$

$$= \int_0^{48.50}\int_0^{19.51} \frac{1}{3}\rho^3 \sin\phi \Big|_{\rho=0}^{\rho=33.17} d\theta\ d\phi$$

$$= \int_0^{48.50}\int_0^{19.51} \frac{(33.17)^3}{3}\sin\phi\ d\theta\ d\phi$$

$$= \int_0^{48.50} \left[\frac{(33.17)^3}{3}\sin\phi\right]\theta \Big|_{\theta=0}^{\theta=19.51} d\phi$$

$$= \int_0^{48.50} 19.51\left[\frac{(33.17)^3}{3}\sin\phi\right] d\phi \qquad \text{where, } \int \sin\phi = -\cos\phi$$

$$= -19.51\left[\frac{(33.17)^3}{3}\right]\cos\phi \Big|_{\phi=0}^{\phi=48.50} \qquad \text{where, } \cos 0 = 1$$

$$= .663\left[-19.51\left[\frac{(33.17)^3}{3}\right]\right] - 1\left[-19.51\left[\frac{(33.17)^3}{3}\right]\right]$$

$$= -157{,}357 - \left[-237{,}341\right] = 79{,}984 \qquad \text{where, multipl. x } \frac{\pi}{180}$$

corrects for angular integrand of $\phi$

$$\therefore \iiint_V dV = 1{,}395.98\,Å^3$$

**4OZF-Vβ**

$\lambda = 120.69°$ $\quad \rho = 23.16\,Å$

$\phi = 180 - 120.69 = 59.31°$ $\quad r = \sin\phi\ (\rho) = 0.860(23.16) = 19.92\,Å$

$\pi \cdot r/4 = 3.14\,(19.92)/4 = 15.64\,Å$

$$\iiint_V dV = \int_0^{59.31}\int_0^{15.64}\int_0^{23.16} \rho^2 \sin\phi\ d\rho\ d\theta\ d\phi$$

$$= \int_0^{59.31}\int_0^{15.64} \frac{1}{3}\rho^3 \sin\phi \Big|_{\rho=0}^{\rho=23.16} d\theta\ d\phi$$

$$= \int_0^{59.31}\int_0^{15.64} \frac{(23.16)^3}{3}\sin\phi\ d\theta\ d\phi$$

$$= \int_0^{59.31} \left[\frac{(23.16)^3}{3}\sin\phi\right]\theta \Big|_{\theta=0}^{\theta=15.64} d\phi$$

$$= \int_0^{59.31} 15.64\left[\frac{(23.16)^3}{3}\sin\phi\right] d\phi \qquad \text{where, } \int \sin\phi = -\cos\phi$$

$$= -15.64\left[\frac{(23.16)^3}{3}\right]\cos\phi \Big|_{\phi=0}^{\phi=59.31} \qquad \text{where, } \cos 0 = 1$$

$$= .510\left[-15.64\left[\frac{(23.16)^3}{3}\right]\right] - 1\left[-15.64\left[\frac{(23.16)^3}{3}\right]\right]$$

$$= -33{,}029 - \left[-64{,}764\right] = 31{,}735 \qquad \text{where, multipl. x } \frac{\pi}{180}$$

corrects for angular integrand of $\phi$

$$\therefore \iiint_V dV = 553.87\,Å^3$$

**4OZG-Vα**

$\lambda = 124.99°$ $\quad \rho = 31.91\,Å$

$\phi = 180 - 124.99 = 55.01°$ $\quad r = \sin\phi\ (\rho) = 0.819(31.91) = 26.14\,Å$

$\pi \cdot r/4 = 3.14\,(26.14)/4 = 20.53\,Å$

$$\iiint_V dV = \int_0^{55.01}\int_0^{20.53}\int_0^{31.91} \rho^2 \sin\phi\ d\rho\ d\theta\ d\phi$$

$$= \int_0^{55.01}\int_0^{20.53} \frac{1}{3}\rho^3 \sin\phi \Big|_{\rho=0}^{\rho=31.91} d\theta\ d\phi$$

$$= \int_0^{55.01}\int_0^{20.53} \frac{(31.91)^3}{3}\sin\phi\ d\theta\ d\phi$$

$$= \int_0^{55.01} \left[\frac{(31.91)^3}{3}\sin\phi\right]\theta \Big|_{\theta=0}^{\theta=20.53} d\phi$$

$$= \int_0^{55.01} 20.53\left[\frac{(31.91)^3}{3}\sin\phi\right] d\phi \qquad \text{where, } \int \sin\phi = -\cos\phi$$

$$= -20.53\left[\frac{(31.91)^3}{3}\right]\cos\phi \Big|_{\phi=0}^{\phi=55.01} \qquad \text{where, } \cos 0 = 1$$

$$= .573\left[-20.53\left[\frac{(31.91)^3}{3}\right]\right] - 1\left[-20.53\left[\frac{(31.91)^3}{3}\right]\right]$$

$$= -127{,}410 - \left[-222{,}356\right] = 94{,}946 \qquad \text{where, multipl. x } \frac{\pi}{180}$$

corrects for angular integrand of $\phi$

$$\therefore \iiint_V dV = 1{,}657.12\,Å^3$$

**4OZG-Vβ**

$\lambda = 128.29°$ $\quad \rho = 21.22\,Å$

$\phi = 180 - 128.29 = 51.71°$ $\quad r = \sin\phi\ (\rho) = 0.875(21.22) = 16.66\,Å$

$\pi \cdot r/4 = 3.14\,(16.66)/4 = 13.08\,Å$

$$\iiint_V dV = \int_0^{51.71}\int_0^{13.08}\int_0^{21.22} \rho^2 \sin\phi\ d\rho\ d\theta\ d\phi$$

$$= \int_0^{51.71}\int_0^{13.08} \frac{1}{3}\rho^3 \sin\phi \Big|_{\rho=0}^{\rho=21.22} d\theta\ d\phi$$

$$= \int_0^{51.71}\int_0^{13.08} \frac{(21.22)^3}{3}\sin\phi\ d\theta\ d\phi$$

$$= \int_0^{51.71} \left[\frac{(21.22)^3}{3}\sin\phi\right]\theta \Big|_{\theta=0}^{\theta=13.08} d\phi$$

$$= \int_0^{51.71} 13.08\left[\frac{(21.22)^3}{3}\sin\phi\right] d\phi \qquad \text{where, } \int \sin\phi = -\cos\phi$$

$$= -13.08\left[\frac{(21.22)^3}{3}\right]\cos\phi \Big|_{\phi=0}^{\phi=51.71} \qquad \text{where, } \cos 0 = 1$$

$$= .620\left[-13.08\left[\frac{(21.22)^3}{3}\right]\right] - 1\left[-13.08\left[\frac{(21.22)^3}{3}\right]\right]$$

$$= -25{,}829 - \left[-41{,}660\right] = 15{,}830.60 \qquad \text{where, multipl. x } \frac{\pi}{180}$$

corrects for angular integrand of $\phi$

$$\therefore \iiint_V dV = 276.30\,Å^3$$

**4OZH-Vα**

$\lambda = 121.57°$ $\quad\rho = 32.35\,Å$

$\phi = 180 - 121.57 = 58.43°$ $\quad r = \sin\phi\,(\rho) = 0.852(32.35) = 27.56\,Å$

$\pi \cdot r/4 = 3.1\bar{4}\,(27.56)/4 = 21.65\,Å$

$$\iiint_V dV = \int_0^{58.43}\int_0^{21.65}\int_0^{32.35} \rho^2 \sin\phi \; d\rho \; d\theta \; d\phi$$

$$= \int_0^{58.43}\int_0^{21.65} \frac{1}{3}\rho^3 \sin\phi \Big|_{\rho=0}^{\rho=32.35} d\theta \; d\phi$$

$$= \int_0^{58.43}\int_0^{21.65} \frac{(32.35)^3}{3}\sin\phi \; d\theta \; d\phi$$

$$= \int_0^{58.43} \left[\frac{(32.35)^3}{3}\sin\phi\right] \theta \Big|_{\theta=0}^{\theta=21.65} d\phi$$

$$= \int_0^{58.43} 21.65 \left[\frac{(32.35)^3}{3}\sin\phi\right] d\phi \qquad \text{where, } \int \sin\phi = -\cos\phi$$

$$= -21.65 \left[\frac{(32.35)^3}{3}\right] \cos\phi \Big|_{\phi=0}^{\phi=58.43} \qquad \text{where, } \cos 0 = 1$$

$$= .524\left[-21.65\left[\frac{(32.35)^3}{3}\right]\right] - 1\left[-21.65\left[\frac{(32.35)^3}{3}\right]\right]$$

$$= -128{,}024 - \left[-244{,}320\right] = 116{,}296 \qquad \text{where, multipl. x } \frac{\pi}{180} \text{ corrects for angular integrand of } \phi$$

$$\therefore \iiint_V dV = 2{,}029.75\,Å^3$$

**4OZH-Vβ**

$\lambda = 130.19°$ $\quad\rho = 21.47\,Å$

$\phi = 180 - 130.19 = 49.81°$ $\quad r = \sin\phi\,(\rho) = 0.764(21.47) = 16.40\,Å$

$\pi \cdot r/4 = 3.1\bar{4}\,(16.40)/4 = 12.88\,Å$

$$\iiint_V dV = \int_0^{49.81}\int_0^{12.88}\int_0^{21.47} \rho^2 \sin\phi \; d\rho \; d\theta \; d\phi$$

$$= \int_0^{49.81}\int_0^{12.88} \frac{1}{3}\rho^3 \sin\phi \Big|_{\rho=0}^{\rho=21.47} d\theta \; d\phi$$

$$= \int_0^{49.81}\int_0^{12.88} \frac{(21.47)^3}{3}\sin\phi \; d\theta \; d\phi$$

$$= \int_0^{49.81} \left[\frac{(21.47)^3}{3}\sin\phi\right] \theta \Big|_{\theta=0}^{\theta=12.88} d\phi$$

$$= \int_0^{49.81} 12.88 \left[\frac{(21.47)^3}{3}\sin\phi\right] d\phi \qquad \text{where, } \int \sin\phi = -\cos\phi$$

$$= -12.88 \left[\frac{(21.47)^3}{3}\right] \cos\phi \Big|_{\phi=0}^{\phi=49.81} \qquad \text{where, } \cos 0 = 1$$

$$= .645\left[-12.88\left[\frac{(21.47)^3}{3}\right]\right] - 1\left[-12.88\left[\frac{(21.47)^3}{3}\right]\right]$$

$$= -27{,}791 - \left[-43{,}087\right] = 15{,}296 \qquad \text{where, multipl. x } \frac{\pi}{180} \text{ corrects for angular integrand of } \phi$$

$$\therefore \iiint_V dV = 266.96\,Å^3$$

**4OZI-Vα**

$\lambda = 125.38°$ $\quad\rho = 31.72\,Å$

$\phi = 180 - 125.38 = 54.62°$ $\quad r = \sin\phi\,(\rho) = 0.815(31.72) = 25.86\,Å$

$\pi \cdot r/4 = 3.1\bar{4}\,(25.86)/4 = 20.31\,Å$

$$\iiint_V dV = \int_0^{54.62}\int_0^{20.31}\int_0^{31.72} \rho^2 \sin\phi \; d\rho \; d\theta \; d\phi$$

$$= \int_0^{54.62}\int_0^{20.31} \frac{1}{3}\rho^3 \sin\phi \Big|_{\rho=0}^{\rho=31.72} d\theta \; d\phi$$

$$= \int_0^{54.62}\int_0^{20.31} \frac{(31.72)^3}{3}\sin\phi \; d\theta \; d\phi$$

$$= \int_0^{54.62} \left[\frac{(31.72)^3}{3}\sin\phi\right] \theta \Big|_{\theta=0}^{\theta=20.31} d\phi$$

$$= \int_0^{54.62} 20.31 \left[\frac{(31.72)^3}{3}\sin\phi\right] d\phi \qquad \text{where, } \int \sin\phi = -\cos\phi$$

$$= -20.31 \left[\frac{(31.72)^3}{3}\right] \cos\phi \Big|_{\phi=0}^{\phi=54.62} \qquad \text{where, } \cos 0 = 1$$

$$= .579\left[-20.31\left[\frac{(31.72)^3}{3}\right]\right] - 1\left[-20.31\left[\frac{(31.72)^3}{3}\right]\right]$$

$$= -125{,}103 - \left[-216{,}067\right] = 90{,}964 \qquad \text{where, multipl. x } \frac{\pi}{180} \text{ corrects for angular integrand of } \phi$$

$$\therefore \iiint_V dV = 1{,}587.63\,Å^3$$

**4OZI-Vβ**

$\lambda = 116.06°$ $\quad\rho = 22.10\,Å$

$\phi = 180 - 116.06 = 63.94°$ $\quad r = \sin\phi\,(\rho) = 0.898(22.10) = 19.85\,Å$

$\pi \cdot r/4 = 3.1\bar{4}\,(19.85)/4 = 15.59\,Å$

$$\iiint_V dV = \int_0^{63.94}\int_0^{15.59}\int_0^{22.10} \rho^2 \sin\phi \; d\rho \; d\theta \; d\phi$$

$$= \int_0^{63.94}\int_0^{15.59} \frac{1}{3}\rho^3 \sin\phi \Big|_{\rho=0}^{\rho=22.10} d\theta \; d\phi$$

$$= \int_0^{63.94}\int_0^{15.59} \frac{(22.10)^3}{3}\sin\phi \; d\theta \; d\phi$$

$$= \int_0^{63.94} \left[\frac{(22.10)^3}{3}\sin\phi\right] \theta \Big|_{\theta=0}^{\theta=15.59} d\phi$$

$$= \int_0^{63.94} 15.59 \left[\frac{(22.10)^3}{3}\sin\phi\right] d\phi \qquad \text{where, } \int \sin\phi = -\cos\phi$$

$$= -15.59 \left[\frac{(22.10)^3}{3}\right] \cos\phi \Big|_{\phi=0}^{\phi=63.94} \qquad \text{where, } \cos 0 = 1$$

$$= .439\left[-15.59\left[\frac{(22.10)^3}{3}\right]\right] - 1\left[-15.59\left[\frac{(22.10)^3}{3}\right]\right]$$

$$= -24{,}624 - \left[-56{,}092\right] = 31{,}468 \qquad \text{where, multipl. x } \frac{\pi}{180} \text{ corrects for angular integrand of } \phi$$

$$\therefore \iiint_V dV = 549.21\,Å^3$$

4GG6-Vα

$\lambda = 132.70°$ $\rho = 30.87$ Å

$\phi = 180 - 132.70 = 47.30°$ $r = \sin\phi\,(\rho) = 0.735(30.87) = 22.69$ Å

$\pi \cdot r/4 = 3.1\overline{4}\,(22.69)/4 = 17.82$ Å

$$\iiint_V dV = \int_0^{47.30}\int_0^{17.82}\int_0^{30.87} \rho^2 \sin\phi \; d\rho\, d\theta\, d\phi = \int_0^{47.30} 17.82\left[\frac{(30.87)^3}{3}\sin\phi\right] d\phi \quad \text{where, } \int \sin\phi = -\cos\phi$$

$$= \int_0^{47.30}\int_0^{17.82} \frac{1}{3}\rho^3 \sin\phi \Big|_{\rho=0}^{\rho=30.87} d\theta\, d\phi = -17.82\left[\frac{(30.87)^3}{3}\right]\cos\phi \Big|_{\phi=0}^{\phi=47.30} \quad \text{where, } \cos 0 = 1$$

$$= \int_0^{47.30}\int_0^{17.82} \frac{(30.87)^3}{3}\sin\phi \; d\theta\, d\phi = .678\left[-17.82\left[\frac{(30.87)^3}{3}\right]\right] - 1\left[-17.82\left[\frac{(30.87)^3}{3}\right]\right]$$

$$= \int_0^{47.30}\left[\frac{(30.87)^3}{3}\sin\phi\right]\theta \Big|_{\theta=0}^{\theta=17.82} d\phi = -118{,}475 - \left[-174{,}742\right] = 56{,}267 \quad \text{where, multipl. x } \frac{\pi}{180} \text{ corrects for angular integrand of } \phi$$

$$\therefore \iiint_V dV = 982.05 \text{ Å}^3$$

4GG6-Vβ

$\lambda = 124.32°$ $\rho = 25.02$ Å

$\phi = 180 - 124.32 = 55.68°$ $r = \sin\phi\,(\rho) = 0.826\,(25.02) = 20.66$ Å

$\pi \cdot r/4 = 3.1\overline{4}\,(20.66)/4 = 16.23$ Å

$$\iiint_V dV = \int_0^{55.68}\int_0^{16.23}\int_0^{25.02} \rho^2 \sin\phi \; d\rho\, d\theta\, d\phi = \int_0^{55.68} 16.23\left[\frac{(25.02)^3}{3}\sin\phi\right] d\phi \quad \text{where, } \int \sin\phi = -\cos\phi$$

$$= \int_0^{55.68}\int_0^{16.23} \frac{1}{3}\rho^3 \sin\phi \Big|_{\rho=0}^{\rho=25.02} d\theta\, d\phi = -16.23\left[\frac{(25.02)^3}{3}\right]\cos\phi \Big|_{\phi=0}^{\phi=55.68} \quad \text{where, } \cos 0 = 1$$

$$= \int_0^{55.68}\int_0^{16.23} \frac{(25.02)^3}{3}\sin\phi \; d\theta\, d\phi = .564\left[-16.23\left[\frac{(25.02)^3}{3}\right]\right] - 1\left[-16.23\left[\frac{(25.02)^3}{3}\right]\right]$$

$$= \int_0^{55.68}\left[\frac{(25.02)^3}{3}\sin\phi\right]\theta \Big|_{\theta=0}^{\theta=16.23} d\phi = -47{,}790 - \left[-84{,}734\right] = 36{,}944 \quad \text{where, multipl. x } \frac{\pi}{180} \text{ corrects for angular integrand of } \phi$$

$$\therefore \iiint_V dV = 644.79 \text{ Å}^3$$

#### Supplement 1.II. Linear Regression Analysis of Calculated Pitch versus *dV* (all V-domains)

| ϕc | *dV* |
|---|---|
| 30.73 | 1703 |
| 12.73 | 297 |
| 30.42 | 1710 |
| 12.21 | 270 |
| 29 | 1512 |
| 13.11 | 406 |
| 18.17 | 740 |
| 25.85 | 1734 |
| 18.81 | 826 |
| 26.03 | 1747 |
| 19.87 | 855 |
| 22 | 1024 |
| 24.75 | 1782 |
| 19.98 | 1198 |
| 26.11 | 2831 |
| 20.96 | 1281 |
| 20.41 | 746 |
| 14.22 | 542 |
| 19.69 | 690 |
| 18.85 | 693 |
| 15.64 | 800 |
| 18.75 | 718 |
| 15.76 | 746 |
| 22.66 | 918 |
| 14.17 | 465 |
| 22.66 | 918 |
| 14.17 | 465 |
| 26.24 | 1397 |
| 15.84 | 554 |
| 26.82 | 1657 |
| 11.92 | 276 |
| 29.32 | 2030 |
| 11.53 | 267 |
| 28.87 | 1588 |
| 16.55 | 549 |
| 22.37 | 982 |
| 15.39 | 645 |

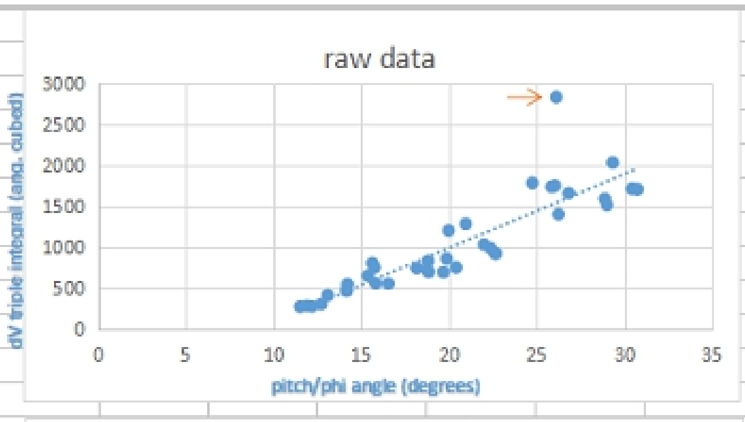


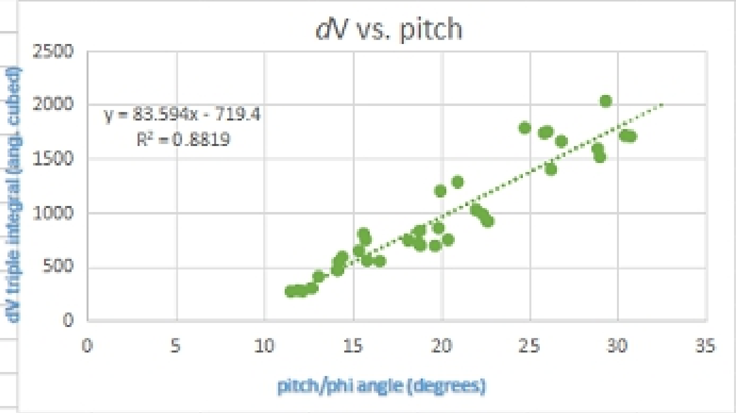


| ϕc | *dV* | *PDB* |
|---|---|---|
| 30.73 | 1703 | 2iam a |
| 12.73 | 297 | 2iam b |
| 30.42 | 1710 | 2ian a |
| 12.21 | 270 | 2ian b |
| 29 | 1512 | 4e41 a |
| 13.11 | 406 | 4e41 b |
| 18.17 | 740 | 1j8h a |
| 25.85 | 1734 | 1j8h b |
| 18.81 | 826 | 1fyt a |
| 26.03 | 1747 | 1fyt b |
| 19.87 | 855 | 3t0e a |
| 22 | 1024 | 3t0e b |
| 24.75 | 1782 | 4h1l a |
| 19.98 | 1198 | 4h1l b |
| → | outlier | 1zgl a |
| 20.96 | 1281 | 1zgl b |
| 20.41 | 746 | 6cql a |
| 14.22 | 542 | 6cql b |
| 19.69 | 690 | 6cqn a |
| 14.45 | 586 | 6cqn b |
| 18.85 | 693 | 6cqq a |
| 15.64 | 800 | 6cqq b |
| 18.75 | 718 | 6cqr a |
| 15.76 | 746 | 6cqr b |
| 22.66 | 918 | 5ksa a |
| 14.17 | 465 | 5ksa b |
| 22.66 | 918 | 5ks9 a |
| 14.17 | 465 | 5ks9 b |
| 26.24 | 1397 | 4ozf a |
| 15.84 | 554 | 4ozf b |
| 26.82 | 1657 | 4ozg a |
| 11.92 | 276 | 4ozg b |
| 29.32 | 2030 | 4ozh a |
| 11.53 | 267 | 4ozh b |
| 28.87 | 1588 | 4ozi a |
| 16.55 | 549 | 4ozi b |
| 22.37 | 982 | 4gg6 a |
| 15.39 | 645 | 4gg6 b |

#### Supplement 1.III. *IMTG* CDR3 Joint Analysis (all V-domains)

**A**

| Chain | V name | 3'V-REGION | N / N1 | D-REGION | N2 | 5'J-REGION | CDR3 | J name | D name |
|---|---|---|---|---|---|---|---|---|---|
| 2iamA | Homsap TRAV22*01 | tg........... | cgcggcgctgattcagggcgcgcagaaactggtg | | | ..........................ttt | C A A L I Q G A Q K L V F | Homsap TRAJ54*01 | |
| 2iamB | Homsap TRBV6-6*01 | tg............... | cgcgagcacctatcat | .........ggc | accggc | ................tatttt | C A S T Y H G T G Y F | Homsap TRBJ1-3*01 | Homsap TRBD1*01 |
| 1FYTA 1J8HA | Homsap TRAV8-4*01 | tg............ | cgcggtgagcgaaagcccgtttggcaacgaaaaactg | | | .........................acctttt | C A V S E S P F G N E K L T F | Homsap TRAJ48*01 | |
| 1FYTB 1J8HB | TRAV28*01 no IMGT junct | tgcgc......gcg agc agc agc acc ggc ctg ccg tat ggc tat acc | | | | .................ttt | C A S S S T G L P Y G Y T F | Homsap TRBJ1-2*01 | no D match |
| 3t0eA | Homsap TRAV26-2*01 | tgca........... | ccgtgtatggcggcgcgaccaacaaactgatt | | | ..............................ttt | C T V Y G G A T N K L I F | Homsap TRAJ32*01 | |
| 3t0eB | Homsap TRBV20-1*01 | tgcag......... | cgcgcgcggcggcagctata | ...acag..... | cccgctgcat | .....................ttt | C S A R G G S Y N S P L H F | Homsap TRBJ1-6*02 | Homsap TRBD1*01 |
| 4h1lA | Homsap TRAV8-3*02 | tg.......... | cgcggtgggcgcgagcggcaacaccggcaaactgatt | | | ...........................ttt | C A V G A S G N T G K L I F | Homsap TRAJ37*01 | |
| 4h1lB | Homsap TRBV19*01 | tg................ | cgcgag | ....cag..... | cctgcgcgatggctataccggcgaa | ...........ctgtttttt | C A S S L R D G Y T G E L F F | Homsap TRBJ2-2*01 | Homsap TRBD1*01 |
| 1zglA | Homsap TRAV9-2*01 | tg............ | cgcgctgagcggcggcgatagcagctataaactgattttt | | | ............................ | C A L S G G D S S Y K L I F | Homsap TRAJ12*01 | |
| 1zglB | Homsap TRBV5-1*01 | tgcgc........... | gagcagcctggcggatcgcgtgaacaccga | ......agcg...... | ttt | ..............ttt | C A S S L A D R V N T E A F F | Homsap TRBJ1-1*01 | Homsap TRBD2*01 |

**B**

| Chain | V name | 3'V-REGION | N / N1 | D-REGION | N2 | 5'J-REGION | CDR3 | J name | D name |
|---|---|---|---|---|---|---|---|---|---|
| 4e41A | Homsap TRAV22*01 | tg.......... | cgcggtggatcgcggcagcaccctgggccgcctgtat | | | ...............................ttt | C A V D R G S T L G R L Y F | Homsap TRAJ18*01 | |
| 4e41B | Homsap TRBV5-8*01 | tg.............. | cgcgag | ....cag..... | ccagattcgcgaaacc | ............cagtatttt | C A S S Q I R E T Q Y F | Homsap TRBJ2-3*01 | Homsap TRBD1*01 |
| 6cqlA 6cqqA 6cqrA | Homsap TRAV24*01 | tg........ | cgcgtttaaagcggcgggcaacaaactgacc | | | ............................ttt | C A F K A A G N K L T F | Homsap TRAJ17*01 | |
| 6cqlB 6cqqB 6cqrB | Homsap TRBV2*01 | tg............... | cgcgagcagccgcctg | .......gcggg.... | cggcatggatgaacagtt | ................tttt | C A S S R L A G G M D E Q F F | Homsap TRBJ2-3*01 | Homsap TRBD1*01 |
| 6cqnA | Homsap TRAV24*01 | tg........ | cgcgtttaaagcggcgggcaacaaactgacc | | | ............................ttt | C A F K A A G N K L T F | Homsap TRAJ17*01 | |
| 6cqnB | Homsap TRBV2*01 | tg............... | cgcgagc | ......agcgg..... | cctggcgggcggcatggatgaacagtt | .................tttt | C A S S G L A G G M D E Q F F | Homsap TRBJ2-3*01 | Homsap TRBD2*01 |
| 5ksaA | Homsap TRAV20*01 | tg........... | cgcggtgcagtttatggatagcaactatcagctgatt | | | .......................tgg | C A V Q F M D S N Y Q L I W | Homsap TRAJ33*01 | |
| 5ksaB | Homsap TRBV9*01 | tg.............. | cgcgagcagcgtg | .......gcggg.... | caccccgagctatgaa | ...........cagtatttt | C A S S V A G T P S Y E Q Y F | Homsap TRBJ2-3*01 | Homsap TRBD2*01 |
| 5ks9A | Homsap TRAV20*01 | tg........... | cgcggtggcgctgaacaacaacgcgggcaacatgctg | | | ........................acctttt | C A V A L N N N A G N M L T F | Homsap TRAJ39*01 | |
| 5ks9B | Homsap TRBV9*01 | tg............. | cgcgagcagcgtggcgcc | ........gggc | agcgatacc | ...........cagtatttt | C A S S V A P G S D T Q Y F | Homsap TRBJ2-3*01 | Homsap TRBD1*01 |

*Genetic calculus of TCR binding class-II pMHC*

| C | V name | Junction | |
|---|---|---|---|
| 4ozfA | Homsap TRAV26-1*01 | 3'V-REGION / N / 5'J-REGION<br>tgcat........... tgcgtttcagggcgcgcagaaactggtg ...........................ttt<br>C I A F Q G A Q K L V F | Homsap TRAJ54*01 J name |
| 4ozfB | Homsap TRBV7-2*01 | 3'V-REGION / N1 / D-REGION / N2 / 5'J-REGION<br>tg............... cgcgagcagctttcgcgcgctg .......gcgg..... cggatacc ............cagtatttt<br>C A S S F R A L A A D T Q Y F | Homsap TRBJ2-3*01 J name<br>Homsap TRBD2*01 D name |
| 4ozgA | Homsap TRAV26-1*01 | 3'V-REGION / N / 5'J-REGION<br>tgcat........... tgtgctgggcggcgcggatggcctg ..............................accttt<br>C I V L G G A D G L T F | Homsap TRAJ54*01 J name |
| 4ozgB | Homsap TRBV7-2*01 | 3'V-REGION / N1 / D-REGION / N2 / 5'J-REGION<br>tg............... cgcgag ....cag..... ctttcgctttaccgatacc ............cagtatttt<br>C A S S F R F T D T Q Y F | Homsap TRBJ2-3*01 J name<br>Homsap TRBD1*01 D name |
| 4ozhA | Homsap TRAV26-1*01 | 3'V-REGION / N / 5'J-REGION<br>tgcat........... tgtgtggggcggcgcgaccaacaaactgatt ................................ttt<br>C I V W G G A T N K L I F | Homsap TRAJ32*01 J name |
| 4ozhB | Homsap TRBV7-2*01 | 3'V-REGION / N1 / D-REGION / N2 / 5'J-REGION<br>tg............... cgcgagc ......agcg...... tgcgcagcaccgatacc ............cagtatttt<br>C A S S V R S T D T Q Y F | Homsap TRBJ2-3*01 J name<br>Homsap TRBD2*01 D name |
| 4oziA | Homsap TRAV4*01 | 3'V-REGION / N / 5'J-REGION<br>tgcct........... ggtgggcgatggcggcagctttagcggcggctataacaaa .......................ctgatttt<br>C L V G D G G S F S G G Y N K L I F | Homsap TRAJ4*01 J name |
| 4oziB | TRAV20*01 no IMGT junct | 3'V-REGION / 5'J-REGION<br>tgc agc.............. gcg ggc gtg ggc ggc cag gaa acc cag tat ........... ttt<br>C S A G V G G Q E T Q Y F | Homsap TRBJ2-5*01 J name<br>no D match D name |
| 4gg6A | Homsap TRAV26-2*01 | 3'V-REGION / N / 5'J-REGION<br>tgcat.......... tctgcgcgatggccgcggcggcgcggatggcctg ...........................accttt<br>C I L R D G R G G A D G L T F | Homsap TRAJ45*01 J name |
| 4gg6B | Homsap TRBV9*01 | 3'V-REGION / N1 / D-REGION / N2 / 5'J-REGION<br>tg.............. cgcgagcagcgtggcggtgagc .......gcggg.... cacctatgaa ............cagtatttt<br>C A S S V A V S A G T Y E Q Y F | Homsap TRBJ2-3*01 J name<br>Homsap TRBD2*01 D name |
